# The GLM-Spectrum: A multilevel framework for spectrum analysis with covariate and confound modelling

**DOI:** 10.1101/2022.11.14.516449

**Authors:** Andrew J Quinn, Lauren Z Atkinson, Chetan Gohil, Oliver Kohl, Jemma Pitt, Catharina Zich, Anna C Nobre, Mark W Woolrich

**Affiliations:** Oxford Centre for Human Brain Activity, Wellcome Centre for Integrative Neuroimaging, University Department of Psychiatry, Warneford Hospital, Oxford, UK; Centre for Human Brain Health, School of Psychology, University of Birmingham, UK; Department for Clinical and Movement Neurosciences, UCL Queen Square Institute of Neurology, 33 Queen Square, London, UK; FMRIB, Wellcome Centre for Integrative Neuroimaging, Nuffield Department of Clinical Neurosciences, University of Oxford, UK; Department of Experimental Psychology, University of Oxford, UK

## Abstract

The frequency spectrum is a central method for representing the dynamics within electrophysiological data. Some widely used spectrum estimators make use of averaging across time segments to reduce noise in the final spectrum. The core of this approach has not changed substantially since the 1960s, though many advances in the field of regression modelling and statistics have been made during this time. Here, we propose a new approach, the General Linear Model (GLM) Spectrum, which reframes time averaged spectral estimation as multiple regression. This brings several benefits, including the ability to do confound modelling, hierarchical modelling and significance testing via non-parametric statistics.

We apply the approach to a dataset of EEG recordings of participants who alternate between eyes-open and eyes-closed resting state. The GLM-Spectrum can model both conditions, quantify their differences, and perform denoising through confound regression in a single step. This application is scaled up from a single channel to a whole head recording and, finally, applied to quantify age differences across a large group-level dataset. We show that the GLM-Spectrum lends itself to rigorous modelling of within- and between-subject contrasts as well as their interactions, and that the use of model-projected spectra provides an intuitive visualisation. The GLM-Spectrum is a flexible framework for robust multi-level analysis of power spectra, with adaptive covariance and confound modelling.

## 1 Introduction

Frequency-domain analyses of oscillations in electrophysiological recordings of brain activity contain information about the underlying neuronal activity. Both the peaks of specific oscillations and the broader spectral shape are informative about brain function and have inspired a wide literature across neuroscience (Buzsakḱi and Draguhn, 2004; Kopell et al., 2014). The time-averaged periodogram is the predominant method for spectrum estimation in neuroscience. It computes the average Fourier spectrum across a set of sliding window segments (Bartlett, 1948, 1950; Welch, 1967) based on the premise that the data are comparable over time and that the effect of noise will be attenuated when averaging across segments. This algorithm produces a statistical estimate of a spectrum and has remained largely the same for many decades. Statistical methods have greatly progressed in this time and many newer approaches can be directly applied to the windowed periodogram.

Here, we propose the General Linear Model Spectrum (GLM-Spectrum) framework for analysing time-averaged periodogram estimates of frequency spectra. This reframes the method of averaged periodograms as a regression problem by modelling frequency spectra over successive windows as a linear mixture of a set of user-specified regressors. This links linear spectrum estimation to the GLM analyses that have been developed for a broad range of neuroimaging applications including structural and functional MRI (Friston et al., 1994; Woolrich et al., 2009), event related fields (Smith and Kutas, 2014) and induced responses (Litvak et al., 2013). Specifically, we demonstrate the utility of multi-level models (Friston, 2007; Woolrich et al., 2004), non-parametric permutation testing (Nichols and Holmes, 2001; Winkler et al., 2014), contrast coding and confound regression in the context of spectrum estimation. GLM-Spectrum can be applied to analyse any time series, from Local Field Potentials to multichannel Electro- and Magnetoencephalography (EEG & MEG). It is not dependent on any specific preprocessing methods in sensor- or source-space analyses, beyond what would apply to a typical frequency spectrum analysis. The GLM-Spectrum could also be configured to perform task analyses on the timescale of the sliding windows used in the STFT. For example, a set of ‘boxcar’ regressors could be defined to contrast different task states. More broadly, the method is applicable to any time series analysis that looks to estimate a Fourier based frequency spectrum.

We illustrate the GLM-Spectrum by analysing EEG recordings alternating between eyes-open and eyes-closed resting-state conditions from a freely available dataset (Babayan et al., 2019). First, the GLM-Spectrum is used to analyse time-series data from a single channel of one individual. The spectrum for the two resting conditions and their difference are computed, whilst a set of covariate and confound regressors account for linear trends over time and a diverse set of potential artefact sources. This approach is generalised to the whole head recording of a single subject to describe the spatial patterns associated with each regressor. Finally, a group-level, whole head analysis explores the GLM-Spectra of specific regressors and contrasts before quantifying how they differ between younger and older participants.

## 2 Methods

### 2.1 Time-Averaged Periodogram Estimation

Time-averaged periodogram methods start by estimating a windowed short-time Fourier transform across time series *y* using the windowing function *w*

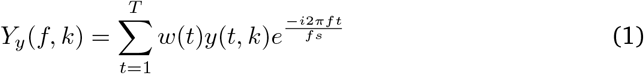

The Fourier transform above is computing the k-th segment of the continuous input *y*(*t*), which we denote with *y*(*t, k*). The output matrix *Y* (*f, k*) contains the STFT, which describes how the spectrum changes in power across the *K* segments. A time-varying magnitude spectrum *S*_*y*_ or power spectrum *P*_*y*_ can be computed from the STFT.

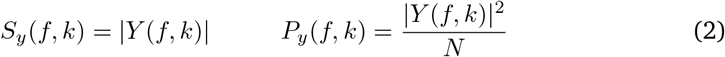

Finally, the time-averaged periodogram is then the average of the time-varying power spectral density across segments.

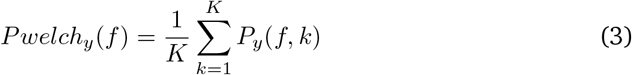

If the previous computations included a windowing function *w*(*t*) and overlapping time segments, then this is Welch’s power spectral density estimate (Welch, 1967). Welch’s time-averaged periodogram now has the property that the noise level of the estimate decreases with increased data length, since more input data provides a larger number of segments for the central averaging step. It is still an imperfect estimator that has been subject to criticism (Prerau et al., 2017; Thomson, 1982) but it is practical, straightforward to compute, and in wide use across science and engineering. A detailed description of these equations is provided in supplemental section 7.1.

### 2.2 General Linear Model Spectrum

The GLM-Spectrum replaces the averaging step in the time-averaged spectrum estimation methods with a General Linear Model (also known as multiple regression). The GLM is widely used in neuroimaging analyses (Friston, 2007; Woolrich et al., 2009) and the same principles around analysis, model validation and statistics apply here. The objective is to model the spectrum across the K sliding window segments as a linear function of a set of regressor variables. The magnitude GLM-Spectrum is defined as:

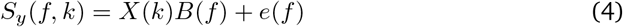

Where *S*_*y*_ (*f*) is the (Kx1) time-varying spectrum estimated at frequency (*f*) across all *K* segments/windows (the STFT computed in 2) computed from a single channel (timeseries) of data, X is a (KxP) design matrix containing the P regressors of interest as they vary over time, and e(f) is a (Kx1) vector of residual errors. We model the whole spectrum using a mass-univariate modelling approach that fits a separate GLM for each frequency bin in the FFT. The resulting (Px1) vector B(f) contains the estimated regression parameters. We refer to the whole vector of estimates across frequency as the GLM ‘beta-spectrum’ ^1^

### 2.3 Estimating the GLM Parameters

Once the design matrix has been specified and the data have been transformed into the STFT, we are ready to fit the regression parameters B in equation 4. Under the assumptions specified above, this can be achieved using Ordinary Least Squares (OLS) to estimate the regression parameters (also known as beta-estimates), *B*(*f*), as

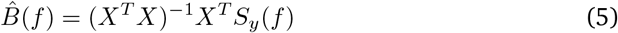

Alternatively, we can pre-multiply the data by the Moore-Penrose pseudo-inverse (MPPI) (Penrose, 1956) of the design matrix, which performs well even when there are multi-collinearities in *X* (see Section 2.4):

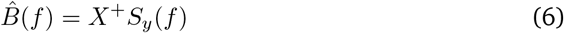

Where the superscript + denotes the MPPI. More complex fitting routines could be used if the assumptions underlying OLS are inappropriate for a particular application. For example, the rest of the GLM-Spectrum framework would work in the same way if *B*(*f*) were estimated using a robust or regularised regression. Similarly, it would be possible to extend the approach to Bayesian regression methods. Here, we use the pseudo-inverse model fitting approach (equation 6) for all GLM estimation.

### 2.4 Assumptions of the GLM-Spectrum

The theory of linear regression underlying the GLM-Spectrum uses assumptions that simplify the problem and specify the conditions under which the solution is valid.Typically, five different assumptions are defined: validity, linearity, independence of errors, homoscedasticity of errors and normality of errors. There remains debate about their relative importance (Gelman and Hill, 2007; Knief and Forstmeier, 2021).

The first two assumptions are relatively general. *Validity* states that the data being analysed should be an appropriate match to the research question. This apparently simple point is frequently overlooked by researchers (Gelman and Hill, 2007). *Linearity* is the assumption that the dependent variable can be described as a linear function of the predictors in the model design matrix. This is the central mathematical feature of linear regression models. We cover assumptions about the residuals and the distribution of variables in more detail in the next two sections.

#### 2.4.1 Distribution of the residuals

Three commonly reported assumptions relate to the residuals of the fitted model *e*(*f*). *Independence of errors* states that the residuals of the model fit are independent and identically distributed (IID) over observations (over time in the case of GLM-Spectrum). *Homoscedasticity of errors* states that the variance of the error is consistent across all values of the predictor. Finally, *normality of errors* states that the residuals should have a normal, Gaussian distribution. Violating these assumptions affects the validity of inferential statistics computed from the model, limiting our ability to generalise results from our data to the population. Parameter estimates are more robust to violations of these assumptions. In most cases, we anticipate that inferential statistics will not be performed on first-level GLM-Spectrum results. Rather the parameter estimates or t-values of many first-level models will be combined into a group model.

GLMs are relatively robust to violations of *homoscedasticity of errors* and *normality of errors* (Williams et al., 2013). The p-values computed from models with violations of these errors tend to be robust at moderate to large sample sizes, except in datasets with substantial outliers (Knief and Forstmeier, 2021).

A specific issue for first-level statistics is the likely presence of temporal autocorrelation in *e*(*f*) indicating a violation of the *independence of errors* assumption. This issue is commonly encountered in other time-series models such as first-level fMRI analyses (Friston et al., 2000; Woolrich et al., 2001; Worsley and Friston, 1995). Future work that requires valid inferential statistics on first-level GLM-Spectra may develop explicit models for this temporal autocorrelation similar to the approach taken in fMRI (Friston et al., 2000; Woolrich et al., 2001).

### 2.4.2 Distribution of the data and predictor variables

The ordinary least squares GLM does not make any formal assumptions about the distributions of the data or predictor variables (Williams et al., 2013). Non-normal predictor variables are commonplace in regression analyses. For example, binary variables that ‘dummy code’ for individual groups of observations are commonly used as predictors. Strongly skewed, fat-tailed or non-normal distributions can still negatively impact the fit by a greater likelihood of influential outlier observations. This may be increasingly problematic with smaller sample sizes, or with increasingly extreme outlier values in the data.

Distribution checking is a critical factor in the choice of whether to use the complex, magnitude, power or log-power spectrum as the dependent variable in a GLM-Spectrum. Power is most commonly used for spectrum analysis but power estimates are strictly positive-valued and tend have distributions with a strong positive skew. The magnitude (*S*_*y*_ (*f*)) and log-power (*log*(*P*_*y*_ (*f*))) spectra are likely to be more Gaussian and both result in similar GLM-Spectrum results (See supplemental section 7.2). Though either of these forms are appropriate, here we use the magnitude spectrum as it is a good compromise between reducing the impact of outliers and maintaining simple visual interpretability of the results.

Future work could equally use the log-power spectrum or consider expanding the GLM-Spectrum approach to use Generalised Linear Modelling (Nelder and Wedderburn, 1972) to account for specific differences in the distribution of the data being modelled. Finally, we do not consider the complex spectrum due to variability in phase across time segments leading to significant cancellation of the signal. If phase information is critical, and expected to be consistent across time segments, then future work may generalise these statistics to the complex spectrum (for example, Baker (2021).

### 2.5 Design Matrix Specification

#### 2.5.1 Regressor Selection

The regressors in the design matrix *X* will typically be secondary time-series that are recorded simultaneously with the main data or known *a priori*. The regressors must be prepared in the same manner as the main data, including any filtering and segmentation, to ensure correspondence between the design matrix and data. All regressors used in this paper are segmented following the modelled time-series data and summed within each segment to create a vector of values to use as a covariate.

The GLM is a highly general method as the design matrix, *X*, can be adapted depending on the application in question. However, this flexibility can also make the specification and interpretation of the regressors challenging. The addition of a new regressor to an existing GLM design matrix can change the parameter estimates and standard errors of the previous regressors. Therefore, the final choice and interpretation of any regressors is necessarily specific to each individual analysis.

Standard time-averaged spectrum estimation methods (such as Welch’s Periodogram) model the mean spectrum across time segments. Similarly, most GLM-Spectrum analyses will also want to include regressors that quantify this average. In the simplest case, a single, constant regressor of ones is directly equivalent to the standard method. However, the flexibility of the GLM allows us to build on this and define more sophisticated models with multiple covariates if required.

One extension enabled by the GLM-Spectrum is to use confound regression to model the effect of an artefact source and attenuate its contribution to the estimate of the overall mean. The amount of denoising applied by the model is proportional to the effect size of the confound regressor in question. This makes the confound regression adaptive to each individual model; the same potential noise source may be highly predictive of the STFT in one dataset but not the next. Researchers could consider running a formal model comparison to remove ineffective confound regressors from first-level analyses altogether. Here, we have taken the approach of maintaining all first-level regressors to simplify group analysis. Further group-level permutation testing the assesses whether a noise source has a ‘significant’ effect on the STFT. This is a flexible alternative to removing the artefactual time periods altogether. Confound regression can be performed by including a non-zero mean regressor alongside a constant regressor in the design matrix. With this specification, the constant regressor models the intercept (the average where the value of the artefact regressor is zero) whilst the confound regressor quantifies the artefact effect. This example is explored in more detail in supplemental section 7.6.

Covariates can be included into the GLM in several ways. We can use indicator regressors (containing zeros and ones) which assume that the covariates effect will be the same each time it is present. Otherwise, we can use dynamic covariates to model phenomena that dynamically change over time in a continuous way. For example, this might include pupil size, heart rate, or respiration rate. When we include these types of continuous regressors, their regression parameters capture the “slope” effect; in other words, how much does the spectrum change with each increment in the value of the regressor. For example, when including a pupil-size regressor, the spectrum resulting from its regression parameter estimates would indicate how much the power in a particular frequency bin increases or decreases as the pupil-size changes by a certain amount.

Another decision is whether to demean a given covariate regressor in the design matrix. Counter intuitively, the interpretation of the regression parameter estimate is unchanged when a covariate is demeaned; in both cases it is modelling the ‘slope’ effect that quantifies how much the spectrum changes with each increment of the regressor. In contrast, the interpretation of a constant regressor in the same model will change depending on whether a covariate is demeaned or not. A constant regressor will model the mean over all time points if the other covariates are demeaned and will model the intercept if non-zero mean regressors are included. As a result, confound regressors that are intended to remove a given effect from the estimate of the mean will typically have a non-zero mean whilst dynamic covariates that model changes around the mean will be demeaned or z-transformed prior to model fitting.

#### 2.5.2 Multicollinearity

Finally, while it is not a violation of the model assumptions, one should take care when regressors in X can be expressed, to any extent, as a linear combination of other regressors. This is referred to as multicollinearity and means that there are infinite equally good solutions to the regression equation. Using the MPPI to fit the model parameters can overcome this limitation. If multiple solutions to equation 4 exist, the MPPI will return the regression parameters with the minimum Euclidean norm (Penrose, 1956). Note that when there is partial multicollinearity, the MPPI uses the component of the regressor that is uncorrelated with the rest of the design matrix (i.e. corresponding to any unique variability in that regressor) to find each parameter estimate. This property means the MPPI solution quantifies the unique effect of each regressor that cannot be accounted for by the others. Therefore, it is frequently desirable to proceed with the MPPI solution for a GLM whose design contains some degree of multicollinearity that we wish to eliminate from the results.

In addition, the impact of any multicollinearity is naturally accounted for in the variance of the affected regression parameter estimates. For example, when the presence of multicollinearity increases uncertainty about the value of a regression parameter then that parameter estimate’s variance (as computed in section 2.6) will be appropriately increased. Nonetheless, even when using the MPPI, we recommend assessing the correlation and singular value spectrum of the design matrix prior to model fitting as well as the variance of the regression parameters (Smith et al., 2007) This ensures that one is aware of the potential impact of multicollinearity on finding a significant result. If these checks identify unexplainable or unintended multicollinearity, perhaps from including too many or inappropriate regressors, then the design should be re-assessed prior to further analysis.

### 2.6 Contrasts and t-statistics

Once the design matrix is specified and the model parameters have been estimated, the GLM-Spectrum consists of a beta-spectrum for each regressor. This beta-spectrum contains the regression parameter estimates quantifying the linear effect of that regressor across the frequency range.

Next, we can compute simple linear combinations of regression parameter estimates, known as contrasts. Contrasts can be defined to ask questions about the size of these linear combinations, including whether they are significantly different to zero (using t-tests). This approach is commonly applied in neuroimaging applications (Friston, 2007; Woolrich et al., 2009).

Each contrast is defined as a vector of values that define the relative weights used to compare different parameter estimates. For example, we could define the following contrasts for a model that contains three regressors in its design matrix:

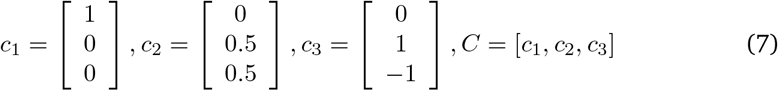

Where C is a (P x *N*_*c*_) matrix containing all *N*_*c*_ contrasts. Using terminology common in neuroimaging, these contrasts define a Contrast Of Parameter Estimates, or a cope, which is computed from a matrix multiplication between the contrast and the model parameter estimates:

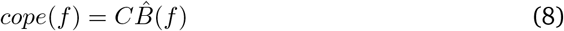

Here, we refer to the resulting frequency resolved vector of cope values, cope(f), as the GLM cope-spectrum. The individual contrasts are designed to ask specific experimental questions. Using the examples in equation 7, the first contrast asks whether 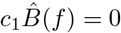. This specifies a t-test that quantifies whether each value in the beta-spectrum of the first regressor is different from zero; regressors two and three are weighted to zero in this specific contrast, but nevertheless still explain variance in the overall model. The second contrast tests if 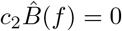 and asks whether the mean of the beta-spectra from regressors two and three is different from zero. Note that setting the values in this contrast to 0.5 ensures that the contrast of the regression parameter estimates can be interpreted as the mean of the two regression parameters involved. When turned into statistics (see below), contrasts *c*_1_ and *c*_2_ are equivalent to one-sample t-tests in classical statistical frameworks.

Finally, testing if 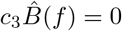tests whether the difference in parameter estimates in the beta-spectrum of regressor 2 minus regressor 3 is different from zero. This is equivalent to an independent samples t-test between the conditions modelled by these regressors. Regressor 1 is set to zero in the second two contrasts and is not directly included in the comparison. However, it is still explaining variance in the model and may be indirectly affecting the outcome of the contrast between regressors 2 and 3.

These contrasts are useful combinations of parameter estimates but we need the associated standard error to complete a formal statistical test. The ratio of the contrast value (cope) and its standard error is a t-statistic that indicates the estimated magnitude of a cope relative to its standard error. To compute the standard errors and subsequent t-statistics for each contrast, we first need to compute the residuals of the model fit:

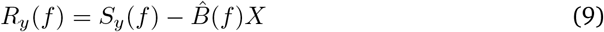

Note that *R*_*y*_ (*f*) contains the actual set of residuals for a given dataset and model fit. This is distinct from *e*_*y*_ (*f*), which denotes a more general white noise process. These residuals are used to compute the variance in the estimate of the cope, also known as a varcope. Firstly, we compute the variance of the residuals:

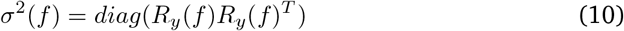

And transform this to get the variance of relevant part of the model for this contrast:

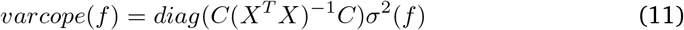

*varcope*(*f*) now contains the square of the standard error for this contrast. This computation can be costly with large datasets as several matrix multiplications must be performed. However, only the diagonal of the resultant matrix is used for further analysis. Therefore, we speed-up this computation using Einstein summation in numpy^2^ to compute only the multiplications which appear in the final diagonal. More information on this and comparisons to alternative computation methods is described in Supplemental section 7.4. The spectrum of t-values corresponding to the contrast can then be computed as the ratio of the cope to its standard error:

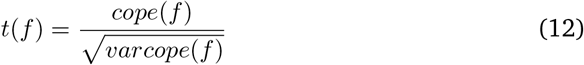

This GLM t-spectrum quantifies the difference of each cope from zero in statistical terms, incorporating both the parameter estimates and their standard errors. Taken together, the GLM beta-spectrum *B*(*f*), cope-spectrum *cope*(*f*) and t-spectrum *t*(*f*) provide an intuitive description of the frequency spectrum of the input data in terms of the specified regressors and contrasts.

### 2.7 Effect Size Computation with Cohen’s *F* 2

Effect sizes represent the strength of a statistical relationship as a complement to hypothesis-based test statistics like t-tests. The effect size of a single variable within the context of a multivariate regression model can be computed with *Cohen*^*′*^*sF* ^2^ (Cohen, 1988). The spectrum of effect sizes for a single regressor within a GLM-Spectrum model can be computed as (Selya et al., 2012).

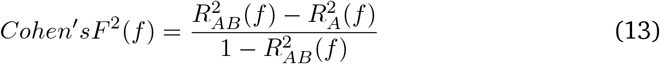

In which *B* denotes the regressor of interest and *A* the set of all other regressors.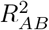 is then the proportion of variance explained by the full model and 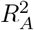 is the variance explained by all regressors except *B*. This metric can be computed for every frequency, channel and regressor within a model to establish a full spectrum of effect sizes. ^3^

Here, we compute effect sizes for each covariate by eliminating it from the full model and computing the value in equation 13. For categorical predictors, the effect size is computed by comparing the full model to a model in which separate categorical regressors are combined into a single constant regressor. As an example, this models the effect of modelling two conditions with their own mean term compared to modelling them both with the same mean.

### 2.8 Visualising effects with model projected spectra

The GLM beta-, cope- and t-spectra assess parts of the overall time-varying spectrum in relation to the model. As these GLM-Spectra can relate to combinations of effects, their impact on the mean spectrum can be difficult to intuit. We propose computing the model-projected spectra to gain a more immediately intuitive visualisation of effects. This is a visualisation of how the spectrum changes for different values of the regressor of interest. For example, if the GLM-Spectrum of EEG data includes a covariate for pupil size then its beta-spectrum will describe how the spectrum changes as pupil size expands and contracts. The model-projected spectrum could then be used to visualise the predicted spectrum at a specific pupil size.

The model-projected spectrum is typically calculated for a descriptive range of values in the original regressor. In this paper, we use the largest and smallest values from the regressor of interest. For example, to compute the projected spectrum for the largest and smallest value of a covariate regressor *Rv* relative to a constant mean term, we use: model-projected spectra

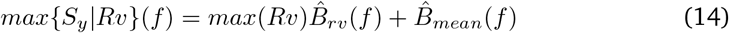

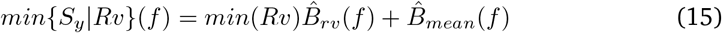

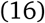

The max and min model-projected spectrum then describes the range of variability in the spectrum that is described by the regressor. Note that this is only a visualisation method and that any apparent differences in these model projected spectra must be confirmed by statistical significance testing, such as non-parametric permutations (see section 2.10).

### 2.9 Group Models for GLM-Spectrum

The GLM-Spectrum described thus far is used to describe continuous data recorded from a single session; we refer to this as the “first-level”. We now consider how we can carry out a “group-level” analysis to combine the results across the first-level GLM-Spectra from multiple sessions/subjects using a group-level (or second-level) GLM (Beckmann et al., 2003; Friston et al., 2002; Woolrich et al., 2004). In brief, we create a group-level dependent variable by concatenating the parameter estimates, copes or varcopes from a set of first-level analyses and use another GLM to model how GLM-Spectra vary over sessions/subjects across the group. For example, here we fit a group-level beta-spectrum using the first-level cope-spectra for N subjects and a group-level design matrix:

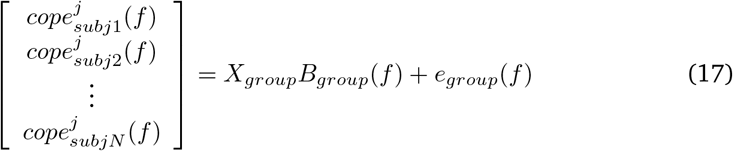

Where 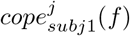 is the *j*^*th*^ first-level cope computed for subject *n* at frequency bin *f*, i.e. 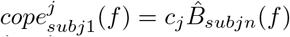where *c*_*j*_ is the *j*^*th*^ first-level contrast and 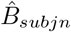is the first-level contrast regression parameter estimates for subject *n*. Note that *X*_*group*_ is the (NxQ) group-level design matrix and the (Qx1) matrix *B*_*group*_(*f*) is the group-level regression parameters, where Q is the number of group-level regressors. As with the first-level GLM, the error *e*_*group*_(*f*) is assumed to be Normal and IID.

**Table 1.** Glossary of definitions for the GLM-Spectrum.

| Term | Definition |
| --- | --- |
| **STFT** | The short-time Fourier transform of a time series, containing the spectrum computed within sliding window time segments across the data. Also called a time-varying spectrum. |
| **Design Matrix** | A matrix of regressors used to explain variability in observed data with a linear regression model. |
| **Regressor** | A single column of a design matrix containing explanatory variables relating to each individual observation. |
| **Beta-estimates** | A parameter estimate describing the linear relation between a regressor and the observed data. Also known as regression parameter estimates. |
| **Beta-Spectrum** | A vector of parameter estimates for a single regressor across the range of a frequency spectrum. |
| **Contrast** | A planned comparison between one or more parameter estimates. |
| **cope** | The result of a defined contrast between beta-estimates. |
| **cope-spectrum** | A vector of cope estimates for a single contrast across the range of a frequency spectrum. |
| **varcope** | The square of the standard error of the cope estimates. |
| **varcope-spectrum** | A vector of estimates for a single contrast across the range of a frequency spectrum. |
| **t-statistic** | The ratio of the departure of the estimated value of a contrast from its hypothesised value to its standard error. |
| **t-spectrum** | A vector of t-statistics for a single contrast across the range of a frequency spectrum. |
| **Model-projected spectrum** | A visualisation of a spectrum as predicted by a fitted GLM set at a particular set of covariate values. |
| **First-Level GLM** | A linear model for a single data recording that models variability across time segments in a STFT with a specified design- and contrast-matrix resulting in a set of beta-, cope- and t-spectra. |
| **Group-Level GLM** | A linear model for a group dataset combining a set of first-level GLM-Spectra. A group-level design- and contrast-matrix models variability in first-level beta or t-spectra across datasets resulting in a set of group-level beta-, cope- and t-spectra. |

As with the first-level analysis, the group-level GLM is fitted using OLS with the MPPI; and is computed separately for each frequency, *f*, (and each channel or voxel – the indexing for which is not shown in the equations) in a mass-univariate manner. In addition, a separate group-level GLM is computed for each first-level cope of interest. As with the first-level GLM, contrasts can be used to ask a range of inference questions from the regression parameter estimates, 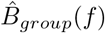. A resource showing examples of commonly used group-level design matrices and contrasts is available online^4^.

As shown in the equation above, the simplest group-level model carries forward the cope-spectra from a set of first-level analyses. This can be thought of as a fixed-effects group model in which each observation (first-level result) contributes equally to the group effect and the group-level GLM models the between-session/subject variability.

This is the approach taken in this manuscript. Future work can extend it to incorporate the information about the first-level standard errors in the varcope-spectra. Both the cope and varcope information could be carried forward to the group-level and fitting a mixed effects model such as the FLAME method in fMRI (Woolrich et al., 2004). In practice, this model is challenging to fit as no simple closed-form estimation is available. Another alternative would be to carry the first-level t-statistics to the group-level. Future work can explore wide range of possibilities for multi-level and mixed-modelling for the GLM-Spectrum.

### 2.10 Non-parametric Permutations for Group GLM-Spectra

Null-hypothesis testing for a given contrast can be carried out with non-parametric permutations (Nichols and Holmes, 2001; Winkler et al., 2014). A null distribution of observed statistics is derived by recomputing the GLM after manipulating the design matrix in line with the null hypothesis. The observed group average is then compared to this null distribution and is ‘significant’ if it exceeds a pre-set critical threshold, such as the 95th percentile of the null distribution. Here, we perform non-parametric permutation using ‘sign-flipping’ for categorical regressors and using ‘row-shuffling’ for parametrically varying regressors.

In all cases, we permute the columns of the design matrix that are directly related to the relevant contrast whilst the remaining regressors are fixed (Draper and Stoneman, 1966; Winkler et al., 2014). The Draper-Stoneman approach can produce erratic results for small sample sizes and may be out-performed by more generally robust methods (Winkler et al., 2014), such as O’Gorman (2005) or Freedman and Lane (1983).

Non-parametric permutation testing could, in principle, be carried out to assess the results of a first-level GLM-Spectrum. In contrast to a group-level analysis, the first-level permutations would only assess whether a particular effect observed within a dataset could be expected to generalise to a wider sample of possible data observations from the same source. Moreover, the time segments in a first-level GLM-Spectrum are likely to exhibit strong autocorrelation (see section 2.4) which would need to be accounted for before any permutation testing could provide a valid result (Friston et al., 2000; Woolrich et al., 2001).

### 2.11 Software implementation and dependencies

The analyses in this paper were carried out in Python 3.9 with core dependencies of numpy (Harris et al., 2020), scipy (Virtanen et al., 2020) and Matplotlib (Hunter, 2007). MNE python (Gramfort, 2013) was used for EEG/MEG data processing with OSL^5^ batch processing tools (Quinn et al., 2022). Python-meegkit^6^ was used for the ‘zapline’ line noise removal algorithm (de Cheveigné, 2020). The spectrum analyses further depend on the Spectrum Analysis in Linear Systems toolbox (Quinn and Hymers, 2020) and glmtools^7^. All code used to run analyses and generate the figures in this paper are available online^8^.

### 2.12 The LEMON Dataset

#### 1.12.1 EEG Preprocessing

All data pre-processing was carried out using MNE-Python and OSL using the OSL batch pre-processing tools. Resting-state EEG recordings from 191 individuals of an open-source dataset (Babayan et al., 2019) were analysed. The raw data for each subject is a resting-state EEG recording from a 62-channel (61 EEG and 1 EOG) using a BrainAmp MR plus amplifier. The channels were in a 10-10 layout and referenced to FCz. Sixteen minutes of data were recorded in one-minute blocks alternating between eyes-closed and eyes-open resting-state. Data were acquired with a bandpass filter between 0.015 Hz and 1 kHz at a sampling frequency of 2500 Hz. The remaining acquisition details are reported in (Babayan et al., 2019).

The raw data were first converted from Brainvision files into MNE-Python Raw data objects. The continuous data were bandpass filtered between 0.25 Hz and 125 Hz using an order-5 Butterworth filter. Line noise was suppressed using a spatial filter following the ‘zapline’ algorithm (de Cheveigné, 2020) as implemented in python-meegkit. Bad channels were automatically identified using the generalised-extreme studentized deviate (G-ESD;Rosner (1983)) routine to identify outliers in the distribution of variance per channel over time. The data were then resampled to 250 Hz to reduce space on-disk and ease subsequent computations.

Independent Component Analysis (ICA) denoising was carried out using a 30 component FastICA decomposition (Hyvarinen, 1999) on the EEG channels. This decomposition explained an average of 99.2% of variance in the sensor data across datasets. Independent components that contained non-neuronal signals such as blinks were automatically identified by correlation with the simultaneous V-EOG channel. ICA components linked to saccades were identified by correlation with a surrogate H-EOG channel, i.e., the difference between channels F7 and F8. Between 2 and 7 ‘artefactual’ components were identified in each dataset, with an average of 2.66 across all datasets. The two ICs that correlated strongest with the V-EOG and H-EOG channels were separately retained for later use in the GLM design matrix.

The continuous sensor data were then reconstructed without the influence of the artefactual V-EOG and H-EOG components. Bad segments were identified by segmenting the ICA-cleaned data into arbitrary 2-second chunks (distinct from the STFT time segments) and using the G-ESD algorithm to identify outlier (bad) samples with high variance across channels. An average of 31 seconds of data (minimum 6 seconds and maximum 114 seconds) were marked as bad in this step. This procedure is biased towards low-frequency artefacts due to the 1/f shape of electrophysiological recordings. Therefore, to identify bad segments with high-frequency content, the same procedure was repeated on the temporal derivative of the ICA-cleaned data. An average of 27 seconds of data (minimum 2 seconds, maximum 109 seconds) were marked as bad when using the differential of the data.

To retain consistent dimensionality across the group, any bad channels were interpolated using a spherical spline interpolation (Perrin et al., 1989) as implemented in MNE-Python. Finally, the spatial gradient between each channel and its distance from the reference sensor (FCz), is attenuated by computing the surface Laplacian (or current source density) of the sensor data. The surface Laplacian data is reference free and has sharper spatial topographies than the raw EEG; though this is a complex computation that is dependent on several hyperparameters, and which may reduce sensitivity to deeper sources (Kayser and Tenke, 2015).

#### 2.12.2 First-Level GLM-Spectrum

A ‘first-level’ analysis models the data within each individual participants EEG data. The STFT is computed for each dataset using a 2-second segment length, a 1-second overlap between segments and a Hanning taper. The 2-second segment length at the sample rate of 250Hz gives a resolution of around 2 frequency bins per unit Hertz in the resulting spectrum. The short-time magnitude spectrum is computed from the complex valued STFT and the frequency bins ranging between 0.1Hz and 100Hz taken forward as the dependent variable in the first-level GLM-Spectrum for that dataset. The GLM design matrix is specified with six regressors (Figure 1). Two binary regressors model intercept term for each of the eyes-open and eyes-closed time segments. The third regressor is a z-transformed covariate describing a linear trend over time. This regressor is included to model potential ‘time on task’ effects over the course of the recording. Previous literature has reported that both alpha power and alpha frequency may change over time within a single EEG recording (Benwell et al., 2019). A non-zero mean regressors for bad segments is computed from the sum of the number of ‘bad’ samples within each STFT time segment. Finally, two further non-zero mean regressors are computed from absolute value of the V-EOG and H-EOG independent component time-courses.

**Figure 1.**
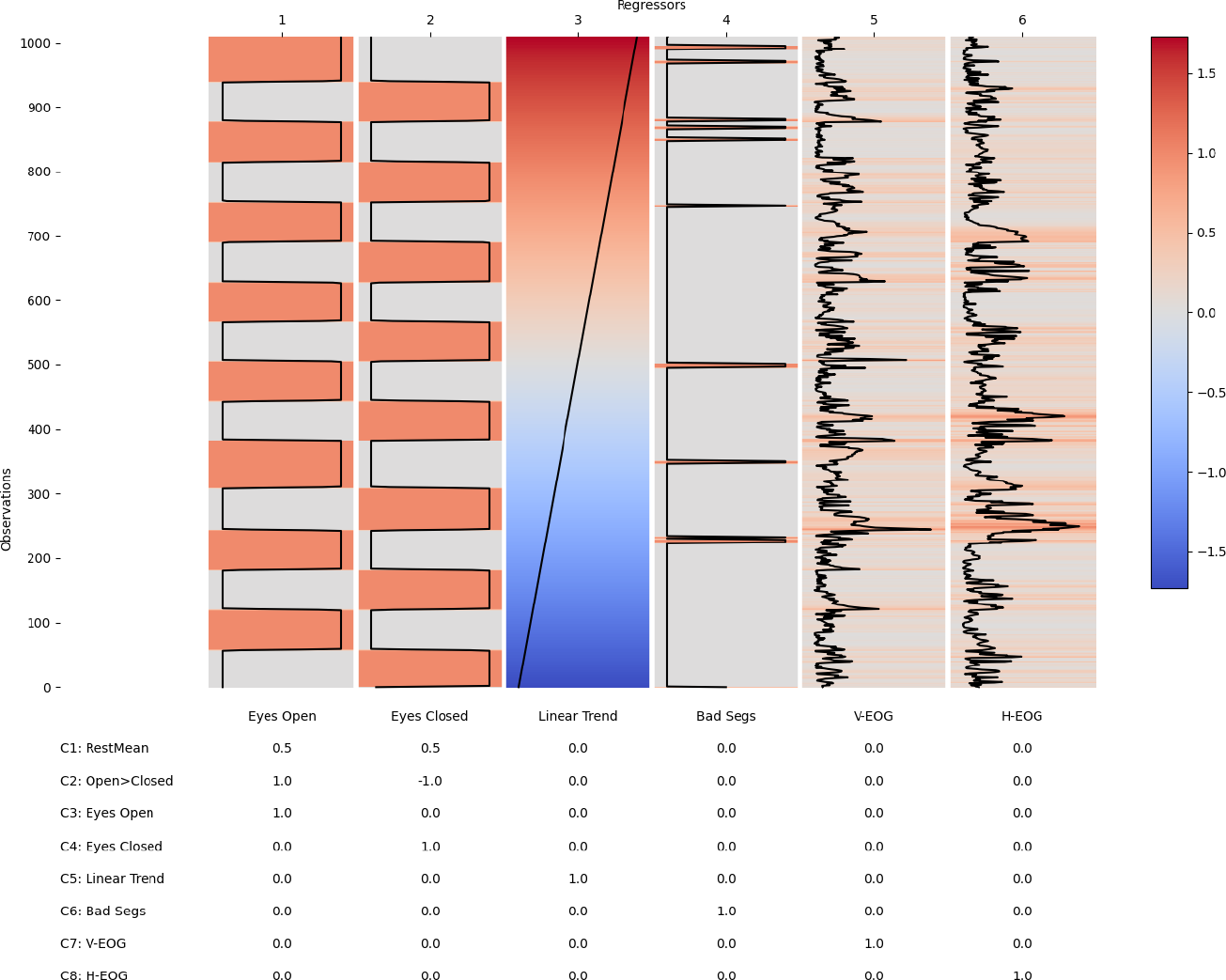
An example first-level design matrix and contrast matrix for a single subject. The top matrix shows the GLM design matrix with individual regressors in columns and single sliding window time segments in rows. The table shows the contrast matrix with corresponding weightings for each regressor. The regressor correlations, singular-value spectrum and variance inflation factors for this design matrix are summarised in Supplemental section 7.5.

A range of contrasts are specified to quantify critical hypothesis tests (see contrast table in Figure 1). The overall mean is modelled by a contrast summing the eyes-open and eyes-closed regressors together weighted by the proportion of ones in each regressor (Contrast 1, Figure 1). A t-test between the spectrum in the eyes-open and eye-closed conditions is specified with a differential contrast (weighted [1, -1], in the direction of eyes-open minus eyes-closed; Contrast 5; Figure 1). Finally, separate

one-sample tests are specified for each covariate with contrasts containing a single 1 for a given regressor (Contrasts 3 to 8; Figure 1). The model parameters were estimated using the MPPI and no statistical assessment was carried out at the first-level.

### 2.12.3 Structural MRI Processing

Individual anatomical details were extracted from T-weighted structural MRI scans (Babayan et al., 2019). All images were processed using the FMRIB Software Library (Woolrich et al., 2009). Images were reoriented to standard Montreal Neurological Institute (MNI) space, cropped, and bias-field corrected. FMRIB’s Linear Registration Tool (FLIRT; Greve and Fischl (2009); Jenkinson et al. (2002); Jenkinson and Smith (2001)) was used to register to standard space before brain extraction was performed using BET (Smith, 2002). Brain images were segmented into different tissue types (grey matter, white matter and CSF) using FMRIB’s Automated Segmentation Tool (FAST; Zhang et al. (2001)). The voxel count for each tissue type was extracted and normalised by the individual’s total brain volume (also computed by FAST) to create a percentage. The total brain volume and individual percentage of grey matter were carried forward as group-level covariates in the GLM-Spectrum.

#### 2.12.4 Group-Level GLM-Spectrum

We now carry out a “group-level” analysis to combine the first-level results and describe between subject variability with another GLM. As described in section 2.12.2, the cope-spectra of each first-level GLM was used as the dependent variable in the group-level GLM. The group-level design matrix contained categorical regressors coding the age group of each participant and covariates corresponding to variability in participant sex, head size and relative grey matter volume (Figure 2). We have modelled age as two distinct, categorical regressors as the LEMON dataset contains participants recruited from either a young (20-40 years old) or an old (60-80 years old) group. Age could equally be included as a single parametrically varying regressor in a dataset that recruited from across a broad age range. A contrast was defined to estimate the mean across all participants and a second to estimate the difference between young and old participants. Finally, one-sample t-tests were specified for each covariate to test whether the regression coefficient significantly differed from zero.

**Figure 2.**
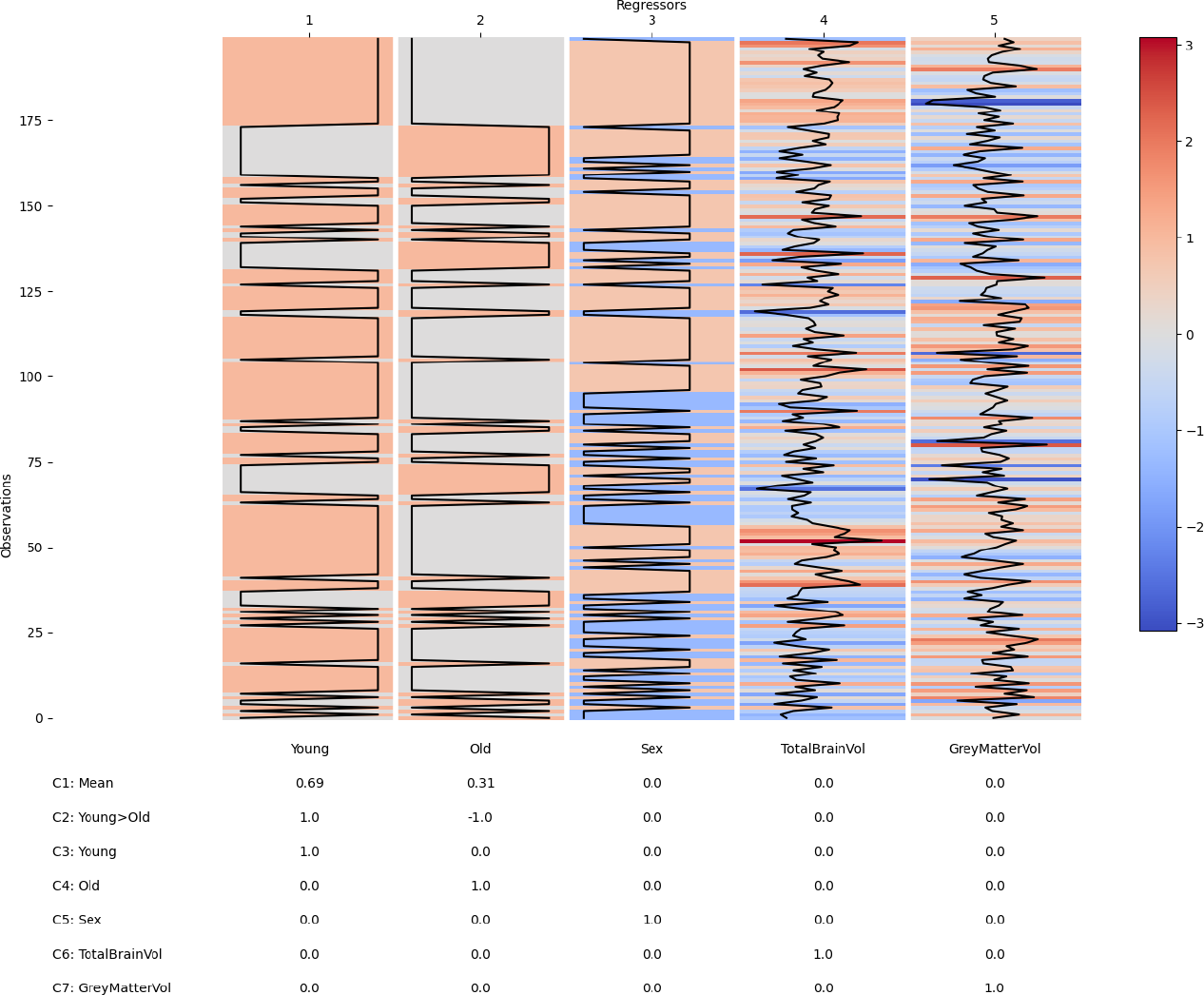
The group-level design matrix and contrast matrix. The matrix shows the design matrix with regressors in columns and individual first-level datasets in columns. The table shows the contrast matrix with corresponding weightings for each regressor. The regressor correlations, singular-value spectrum and variance inflation factors for this design matrix are summarised in Supplemental section 7.5. This group design matrix was used to to model variability across datasets in each of the first-level contrasts.

The group-level design matrix was used to model each of the first-level contrasts. The group model parameter estimates were computed using the MPPI. Statistical significance in the group-level t-spectra was assessed using cluster-based non-parametric permutations using sign-flipping or row-shuffle permutations. A cluster forming threshold of p=0.001 was used (equivalent to t(205)=3.34) across all tests. A spatial extent threshold of 1 was set to ensure that any cluster has at least 2 adjacent points exceeding the cluster forming threshold.

## 3 Results

### 3.1 First-level covariate spectra on a central EEG channel

Figure 2 summarises the first-level GLM-Spectrum analysis of a single channel (Pz) from an exemplar resting-state EEG recording. The pre-processed EEG time series (Figure 3A) was split into 2 second time segments with a 50% overlap (Figure 3B), modified by a Hann window (Figure 3C) and transformed into the frequency domain using a Short-Time Fourier Transform (STFT; Figure 3D). Each column of this STFT contains the time course of the magnitude at each frequency and constitutes the dependent variable in a first-level GLM. The first-level GLM (see methods section 2.12.2) models this variability over time in relation to the resting conditions, artefacts detected in the data and the electrooculogram (EOG). The final first-level design matrix had six regressors (Figure 3E) modelling the two resting-state conditions, a linear trend over time and three potential dynamic confounds. The full design matrix and contrast specification for the entire run can be seen in Figure 1.

**Figure 3.**
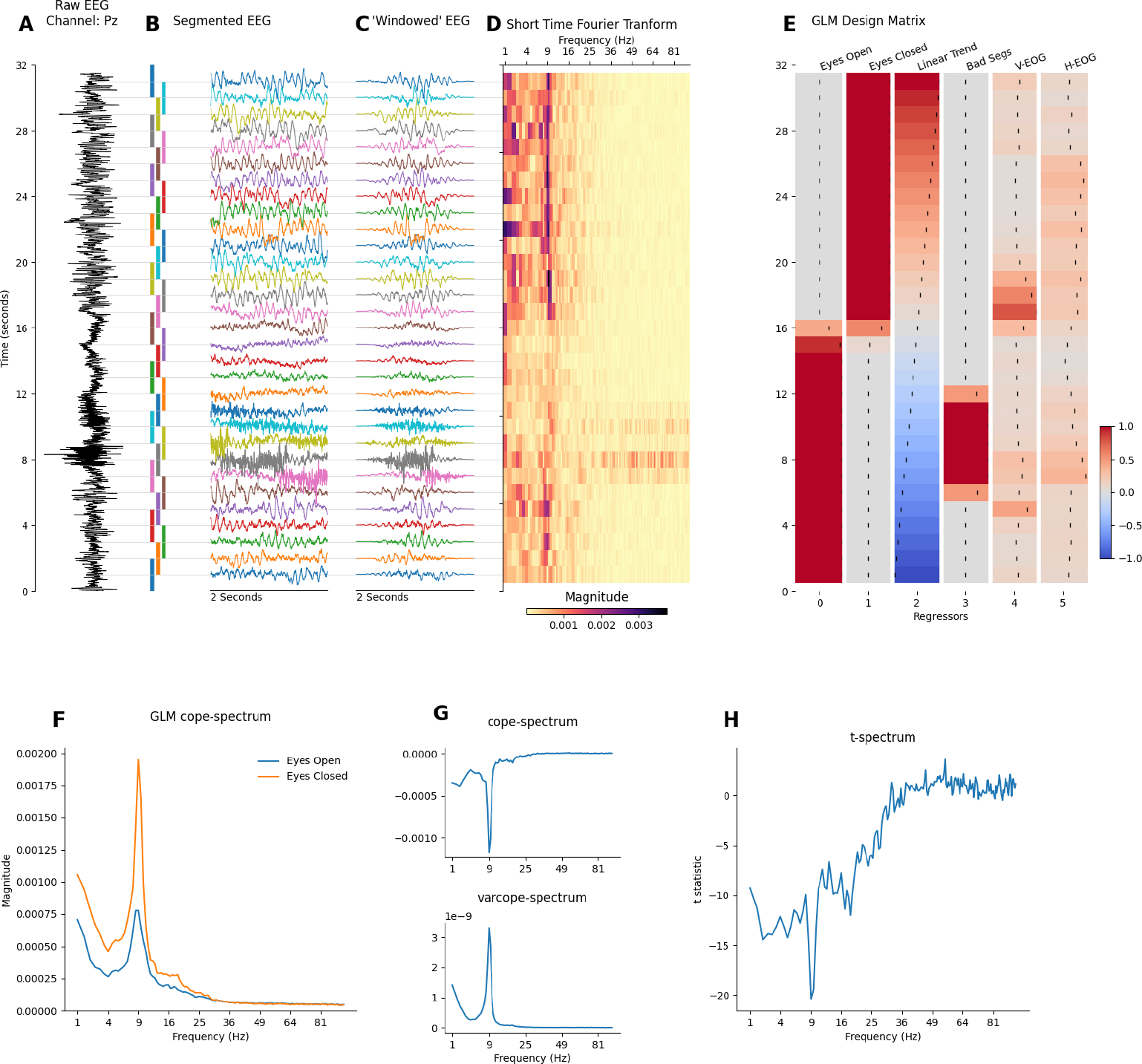
First-level (i.e. within-session) GLM-Spectrum description during alternating eyes-open and eyes-closed resting-state from channel Pz in a single EEG recording. **A:** A 32-second segment of pre-processed EEG time-course from sensor Pz. **B:** EEG time-course segmented into 2-second sliding windows with 50% overlap. **C:** Windowed data segments modified by a tapered Hann window function. **D:** Short time-Fourier transform computed with the FFT of each windowed data segment. Each column of this matrix (change in magnitude of a single frequency over time) is the dependent variable to be described by the GLM. **E:** The GLM design matrix containing condition, covariate and confound regressors. This is zoomed in section of the full design matrix in Figure 1, showing only rows corresponding to the data shown here. **F:** The GLM beta-spectra for the two regressors modelling the spectrum during eyes-open and eyes-closed rest (contrasts 3 and 4 in Figure 1). **G:** The cope- and varcope-spectrum for a differential contrast between the eyes-open and eyes-closed conditions as specified in contrast 5 in Figure 1. **H:** The t-spectrum for the contrast in G.

The first-level GLM was fitted separately for each frequency bin using a standard ordinary least squares routine. The average magnitude spectra for the eyes-open and eyes-closed rest periods are quantified by two cope-spectra (Figure 3F) specified by first-level contrasts 3 and 4 (Figure 1). Both cope-spectra showed a 1/f-type structure and a prominent alpha peak around 9 Hz. The eyes-open > eyes-closed contrast (the regressor was coded to have ones for eyes-open and minus ones for eyes-closed; specified in contrast 5 in Figure 1) had negative values in the cope-spectrum indicating that spectral magnitude was larger in the eyes-closed condition across a range of frequencies, peaking around the alpha range (Figure 3G – top panel). The square of the standard error of the estimates is shown in the varcope-spectrum (Figure 3G – top panel) and indicates where the estimate of the mean was least certain. This roughly followed the shape of the spectra in Figure 3F, showing a clear alpha peak and a weak 1/f trend. This is an example of a close relationship between cope and varcope estimates that can lead to large effects in the beta-spectrum being substantially less prominent in the t-spectrum. The final t-spectrum contained a full spectrum of t-values for the contrast between the two resting conditions (Figure 3H). The large magnitude, negative t-values in alpha and surrounding frequencies qualitatively indicated a greater magnitude in the eyes-closed condition. In sum, this shows a full spectrum perspective on the ‘alpha reactivity’ or ‘alpha blocking’ effect (Adrian and Matthews, 1934) that is most commonly assessed within a-priori frequency bands (Babiloni et al., 2011; Wan et al., 2018).

### 3.2 First-level spectral analysis on whole head EEG

So far GLM-Spectrum method has been applied to univariate data (i.e., single channel EEG data), but it can be readily extended to a full multichannel dataset. To model the GLM-Spectrum across channels, a separate GLM using the same design matrix was fitted to each channel and frequency bin. This provides a description of spectral effects over frequency and space. Accordingly, we extended the resting-state model to the 61-channel whole head EEG recording. The design matrix and contrast specification were the same as the single channel analysis.

The beta-spectrum computed from the condition regressors showed the familiar 1/f slope and prominent occipital alpha features in both resting-state conditions. The GLM analysis was identical to the single channel results (Figure 3) but now can include spatial distributions alongside the frequency spectrum. Qualitatively, the eyes-open condition (Figure 4A) had a smaller alpha peak than the eyes-closed condition (Figure 4B). Both conditions had a similar topography in this subject. The t-spectrum of the contrast between eyes-open and eyes-closed rest showed a large, negative effect peaking around the alpha range (Figure 4C) as seen in the single channel example (Figure 3G). This difference had a spatial maximum in posterior-central regions, replicating the occipital-parietal location of the alpha reactivity effect widely reported in the literature (Babiloni et al., 2011; Wan et al., 2018).

**Figure 4.**
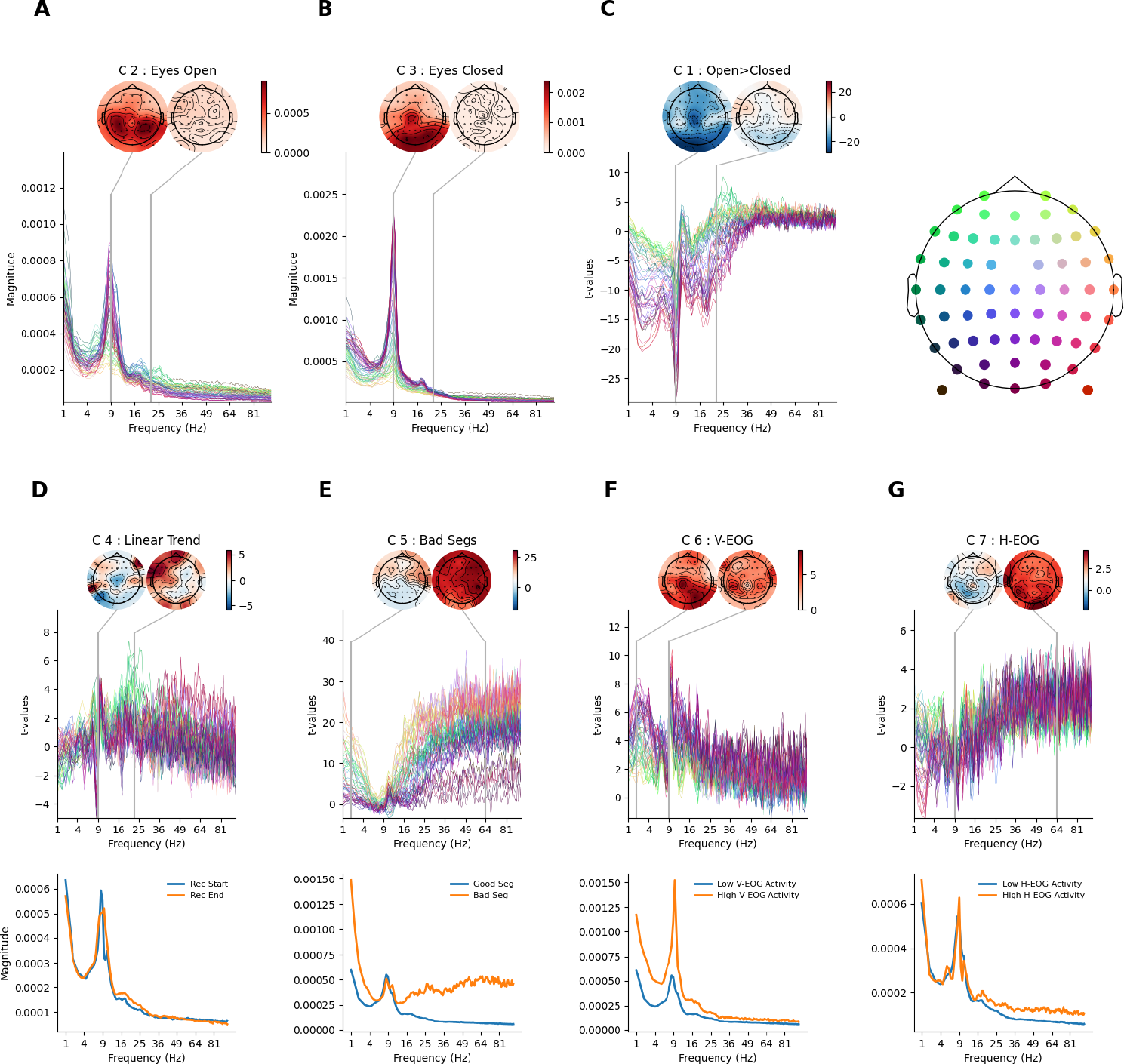
GLM-Spectrum fits for alternating eyes-open and eyes-closed resting-state EEG for a single participant. Mean magnitude spectrum estimates for each channel and frequency bin. The topography (top right) provides location-colour coding used in the t-spectra throughout the figure. The colour range of each topography is set vary between plus and minus the maximum absolute value of the data plotted. The contrasts shown in this figure are detailed in supplemental section H. **A:** Beta-spectrum for the eyes-open condition regressor. The topography shows the spatial distribution of spectral magnitude at 9 Hz **B:** Beta-spectrum for the eyes-closed condition regressor, layout is same as in A. **C:** t-spectrum for the contrast between the eyes-open and eyes-closed conditions. **D:** t-spectrum spectrum for the linear trend covariate (top). Model-predicted magnitude spectra averaged across all sensors for the extrema of the predictor, which in the case of a linear trend regressor corresponds to the start and end of the scan (bottom). **E:** t-spectrum spectrum for the bad segment confound (layout same as for D), model-projected spectra are shown for good and bad segments. **F:** t-spectrum spectrum for V-EOG confound (layout same as for D), model-projected spectra are shown for the maximum and minimum observed V-EOG activity. **G:** t-spectrum spectrum for H-EOG segment confound (layout same as for D), model-projected spectra are shown for the maximum and minimum observed, H-EOG activity.

The linear trend showed substantial t-values around the alpha range, though its model projected spectra suggest that this is a subtle effect (Figure 4D). The bad segment regressor effect (Figure 4E) peaks at relatively low (<4Hz) and high (>25Hz) frequency ranges. The size of this effect is visualised more intuitively in the model projected spectrum (Figure 3F – bottom panel) which showed the substantial increase in both a low- and high-frequency range. Finally, the V-EOG (Figure 4F) and H-EOG (Figure 4G) covariates both showed large t-values in relatively low frequencies (around 1Hz). The V-EOG had a large negative effect around the alpha range, suggesting that time segments with high V-EOG activity were associated with lower alpha magnitude. In contrast, the H-EOG showed an additional positive effect in high frequencies suggesting that segments with high H-EOG activity are associated with larger high frequency spectral magnitude. In particular, the complex pattern of effects in the V-EOG t-spectrum manifests as segments with high V-EOG activity showing a decrease in alpha magnitude and an increase in alpha frequency.

### 3.3 Covariate effects are highly variable across datasets

Next, we build on the single subject exemplar result by exploring the effect of the covariate and confound regressors across 206 EEG recordings from the LEMON dataset. Firstly, we show a qualitative description of the model R-squared values and related effect sizes for each predictor variable. The group averages here are intended as illustrations and are not supported by inferential statistics. Firstly, the GLM-Spectrum was able to describe between 60% and 70% of the variance in the STFT, averaged across all participants and channels (Figure 5A). This value was highly variable across participants, particularly above 50Hz where the R-square values ranges between 20% and 80%. Secondly, Cohen’s *F*^2^ statistic was used to compute the marginal effect size of each predictor within the model (see section 2.7 for details). Modelling the two resting conditions separately, as opposed to using a single average, results in a group average effect size which peaks around the alpha frequency, though the effect in individual datasets was highly variable (Figure 5B). The effect size of the four covariates has a low group average but is, again, highly variable in individual datasets (Figure 5C). This variability was particularly prominent at higher frequencies and indicates the adaptive effects of the covariate regressors. These predictors have strong associations with the STFT in some individuals but not others.

**Figure 5.**
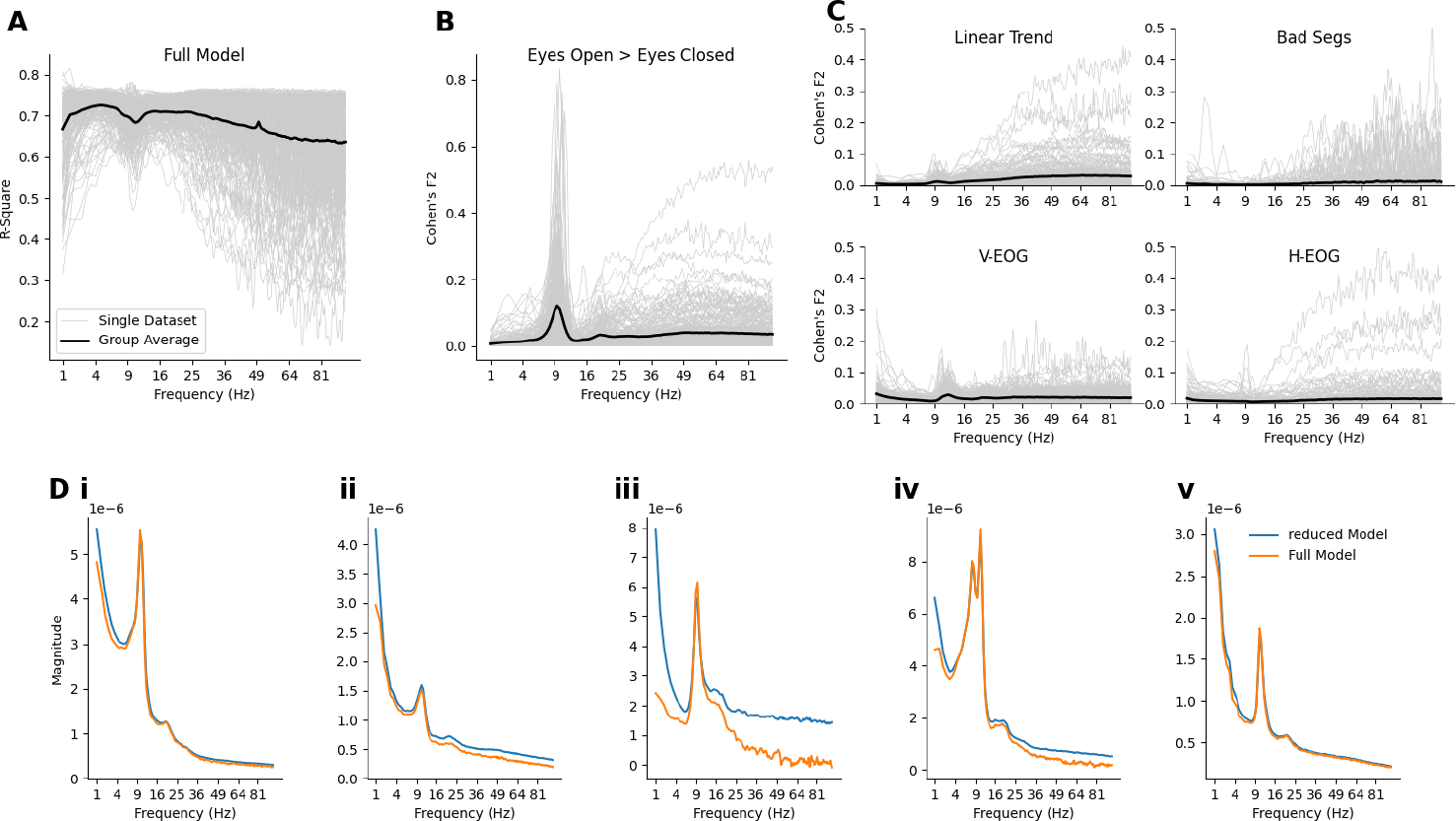
Effect of first-level confound variables on mean spectrum estimates. **A:** The proportion of variance explained by the full model for all datasets. The values are averaged across channels and plotted across frequency. The grey lines represent the effect size spectra of individual datasets and the black line is the group average across all datasets. **B:** Cohen’s *f*^2^ values for the eyes-open *>* eyes-closed contrast. Plotting style is the same as in A). **C:** Cohen’s *f*^2^ values for the each of the four covariates. Plotting style is the same as in A). **D:** Detailed visualisation of the average beta-spectra across all channels from five example datasets.

To further illustrate the effect of including the continuous covariate predictors the frequency spectrum was computed for both the full model and a reduced model containing only the two condition terms (i.e., eyes-open and eyes-closed), excluding other covariates and confounds (Figure 5D). The inclusion of covariate regressors has a different pattern of effect across frequencies for each dataset. The difference between the full and reduced model is relatively small for datasets i and v indicating that the effect of the covariates was minimal for these recordings. In contrast, datasets ii and iv show moderate differences in low and high frequencies whilst iii shows a substantial difference between the full and reduced model.

### 3.4 Group-level design matrix and contrasts

The GLM-Spectrum framework can be extended to group analyses by carrying a set of first-level results to a second-level GLM (See methods section 2.9). This group-level analysis models between-subject variability across independent first-level GLM-Spectra. These multilevel, hierarchical models are well established in neuroimaging (Beckmann et al., 2003; Friston et al., 2002; Friston, 2007; Woolrich et al., 2004), and the existing theory applies to the GLM-Spectrum.

The group-level GLM was fitted to the first-level cope-spectra (Figure 6A) across all datasets separately for each channel, frequency bin and first-level contrast (see methods section 2.12.4). The group-level design matrix contained two condition regressors modelling the mean across subjects for younger and older participants separately and three z-transformed parametric covariates modelling between-subject variability in sex, total brain volume and relative grey matter volume (Figure 6B). Two group contrasts were defined alongside the main effects. One overall average that modelled the sum of the young and old groups (Contrast 1; Figure 6B) weighted by the number of participants in each group. A second contrast quantified the linear group difference (Contrast 2; Figure 6B). Finally, a set of main effect contrasts were also defined for each regressor (Contrasts 3 to 7; Figure 6B). The final fitted model contains a group-level beta-spectrum that describes the linear effect of a group regressor across separate datasets. The group-level GLM returns beta-spectra for the overall mean spectrum (Figure 6C) as well as contrasts and main effects (Figure 6D). At the group-level, the error bars now indicate the standard error of the fitted mean across participants (rather than across STFT time windows, as in the first-level).

**Figure 6.**
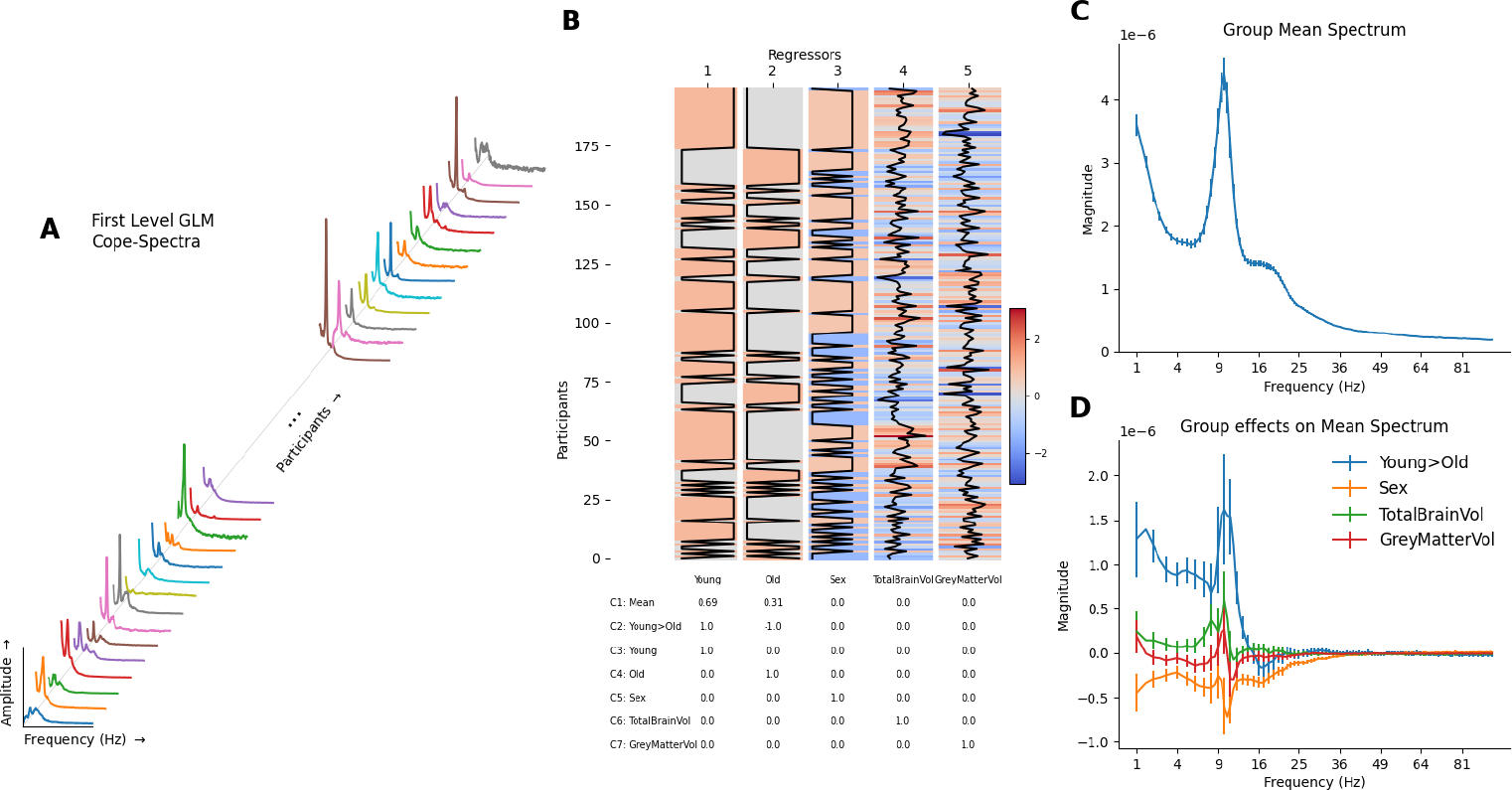
Group-level GLM-Spectrum describing a group mean and variability associated with two between-subject factors. **A:** The data modelled by the group-level GLM are the first-level GLM-Spectra across all participants. A GLM is fit separately for each channel, frequency bin and first-level contrast. **B:** The group-level design matrix. The first and second regressors are categorical predictors coding young and old participants. **C:** The group-level estimate obtained from the first contrast coding for the weighted average of the two regressors (contrast 1). Error bars indicate the varcope of the group contrast. **D:** The group-level effects for the difference between young and old participants (contrast 2, weighted [1, -1]) and the three covariate contrasts (contrasts 5, 6 and 7), quantifying the effect of sex, total brain volume and normalised grey matter volume respectively. Error bars indicate the varcope of the group contrast.

### 3.5 Group effects of age and eyes-open versus eyes-closed rest

Next, we explored how the GLM-Spectrum varies across resting conditions and participant age. Therefore, two main-effect contrasts and their interaction were explored. Two-tailed, non-parametric, cluster (with clusters formed over frequencies and sensors) permutation tests were used to establish statistical significance for all group-level analyses.

We first computed the group average of the within-subject difference between eyes-open > eyes-closed rest. Specifically, we computed the *Mean* group-level contrast (Contrast 1; Figure 6B) on the *eyes-open > eyes-closed* first-level contrast (Contrast 2; Figure 1). Non-parametric cluster permutation testing indicated two significant clusters (Figure 7A) that together cover the whole frequency range investigated here. The first cluster showed a negative effect with greater magnitude in the eyes-closed condition. This cluster spanned the lower frequency range (approximately 30Hz and lower) across almost all channels, though the effect peaked around the alpha range in occipital channels. The posterior peak within this cluster matched the expected occipito-parietal source of the alpha reactivity effect (Wan et al., 2018). The second cluster showed a positive effect indicating higher magnitude in the eyes-open condition. This cluster spanned the higher frequency range (10-100 Hz) across all channels with the largest effect in bilateral frontal regions. This cluster may partially reflect residual eye movements that have not been accounted for by ICA or the first-level artefact regression. The t-statistics in a low frequency range in frontal sensors are strongly modulated by the inclusion or exclusion of the EOG based confound regressors (see supplemental section 7.7).

**Figure 7.**
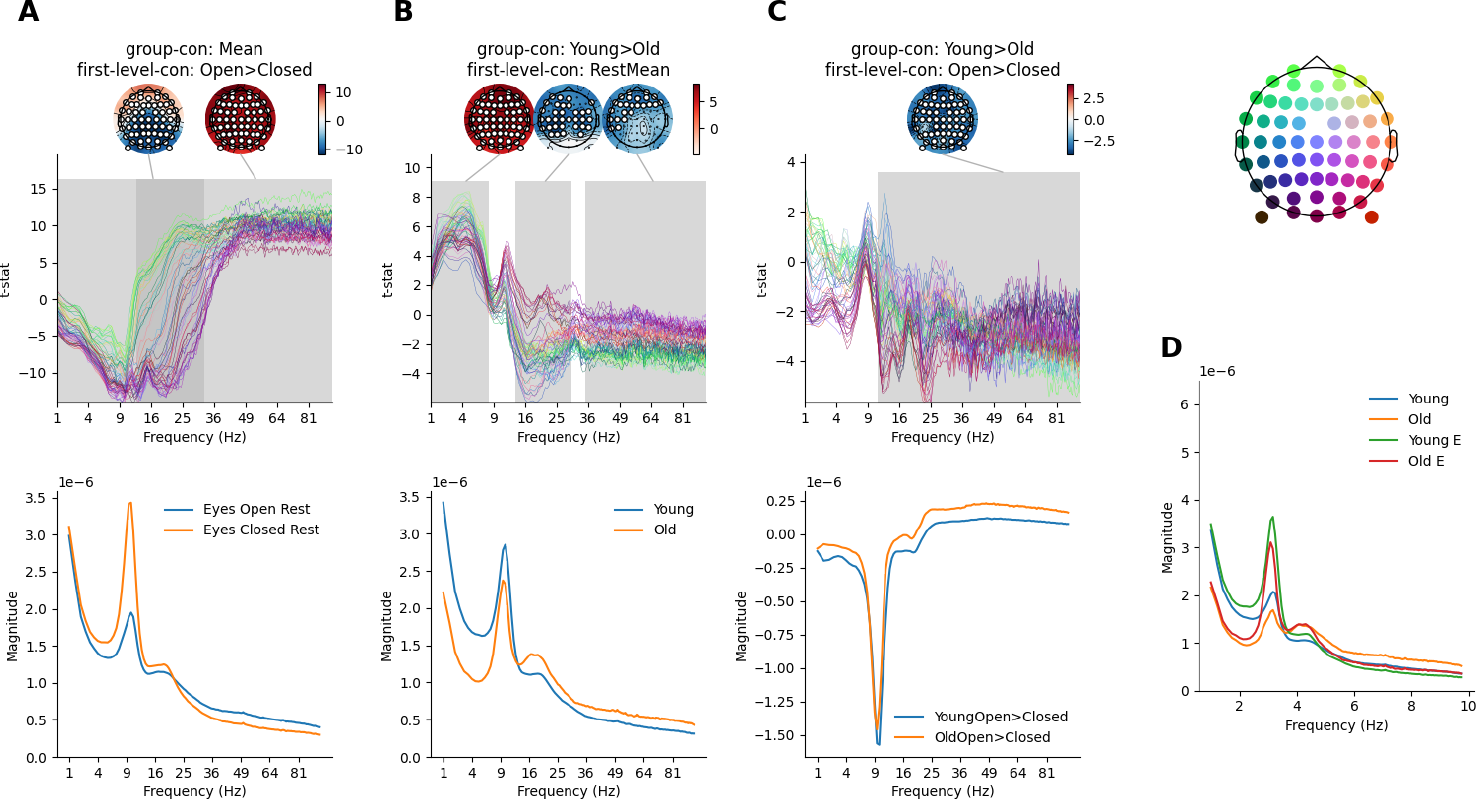
Higher order GLM-Spectrum group-level results. The topography (top right) provides location-colour coding used in the t-spectra throughout the figure. In all cases, statistical significance is assessed by sensors x frequency cluster permutation testing and is indicated in grey. The colour range of each topography is set to vary between plus and minus the maximum absolute value of the data plotted. **A:** The within-subject contrast between eyes-open and eyes-closed conditions, averaged across all participants. Model-predicted magnitude spectra averaged across all sensors for the extrema of the predictor (bottom). **B:** The between-subject difference in average magnitude between young and old participants. Layout is the same as for A. **C:** The interaction effect exploring whether the eyes-open>eyes-closed contrast varies between young and old participants. Layout is the same as for A. **D:** The averaged GLM-Spectra for the two conditions and two groups.

Further, we computed the difference in the time-averaged first-level spectra between the younger and older adults. This corresponds to computing the *Young > Old* group-level contrast (Contrast 2; Figure 6B) on the *Overall Mean* first-level contrast’s cope-spectrum (Contrast 1; Figure 1). Non-parametric permutations on this showed three significant clusters. The first cluster indicated a positive effect in which younger participants had larger power than older participants. The cluster covered low frequencies (<8 Hz) and all channels peaking in frontal and occipital channels (Figure 7B). The direction of this effect is consistent with previously reported decreases in delta and theta power in older adults (Klimesch, 1999). The second cluster indicates greater beta-frequency magnitude for older adults. The cluster spans between 15 and 30Hz and peaks in bilateral central sensors. This finding replicates literature showing higher beta power in older adults (Xifra-Porxas et al., 2019). Finally, the third cluster indicates greater magnitude for older participants in a cluster between 35 and 100Hz that peaks in frontal sensors. In addition, the change in overall spectral shape qualitatively supports indications that older adults have a flatter 1/f slope in the EEG spectrum (Merkin et al., 2022; Voytek et al., 2015), though we did not explicitly quantify 1/f slope here.

No significant cluster for an age difference was identified in the alpha range, though individual t-statistics reach around 5. This null effect may relate to the choice of sensor normalisation during pre-processing (Klimesch, 1999). In addition, the choice of a 0.5 to 100 Hz frequency range may be too wide to assess any relatively narrow band alpha changes in the presence of large broadband effects in higher frequencies.

Finally, we explored whether the within-subject difference in eyes-open and eyes-closed resting-state changed between the younger and older adults. Specifically, this corresponds to the *Young > Old* group-level contrast (Contrast 2; Figure 6B) computed on the *eyes-open > eyes-closed* first-level contrast (Contrast 5; Figure 1). Non-parametric permutation testing identified a single significant cluster with a negative t-statistic (Figure 7C). This indicates the presence of an interaction effect in which older adults show a larger difference in the eyes-open > eyes-closed contrast.

Interestingly, there is no indication of an interaction effect in the low to mid alpha range. The interaction can be qualitatively summarised by plotting the beta-spectrum separately for each of the underlying four mean levels (Figure 7D).

### 3.6 Group average of first-level cope-spectra

We explored the group averages of each first-level covariate regressor. The first-level linear trend regressor is expected to be sensitive to slow drifts throughout the recording. The t-spectrum of the *Mean* group-level contrast (Contrast 1; Figure 6B) computed on the *Linear Trend* first-level contrast (Contrast 5; Figure 1) showed four significant clusters (Figure 81) with a mix of increases and decreases across space and frequency. Broadly these clusters showed a decrease in magnitude over time in both low frequencies (0.5-4Hz) and high frequencies (20-100Hz) peaking in occipital posterior channels. In contrast, an increase in magnitude over time was found in a low theta/alpha range (3-9Hz) and high frequencies (15-100Hz) peaking in frontal sensors. The presence of bad segments was associated with a single, strong positive effect indicating increased magnitude in the EEG during marked bad segments. The *Mean* group-level contrast (Contrast 1; Figure 6B) computed on the *Bad Segments* first-level contrast (Contrast 6; 1) showed a strong positive effect across all frequencies and across all channels (Figure 8B).

**Figure 8.**
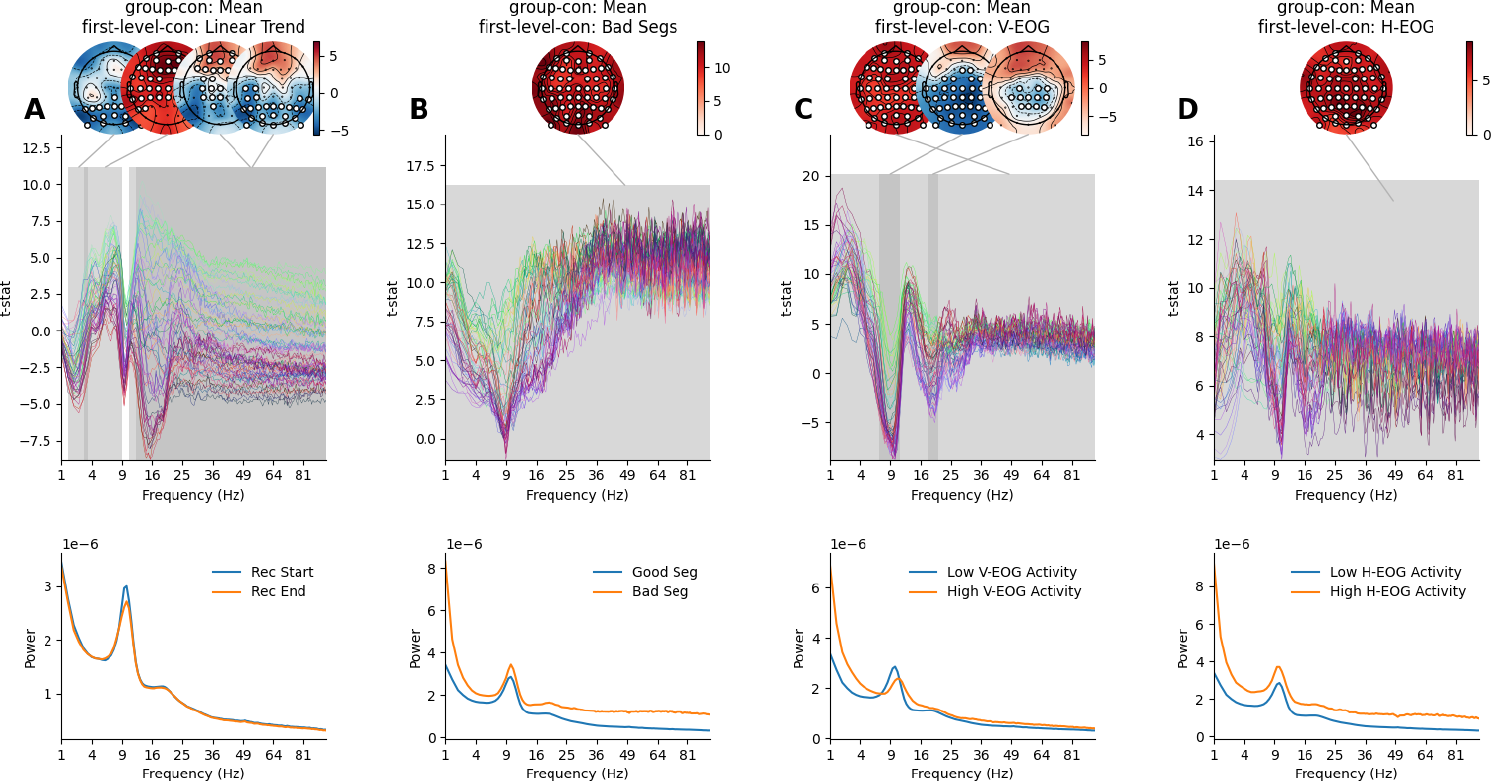
Group-level GLM-Spectrum results for each first-level regressor. The topography (top right) provides location-colour coding used in the t-spectra throughout the figure. In all cases, statistical significance is assessed by sensors x frequency cluster permutation testing and is indicated in grey. Topographies for the largest three significant clusters are shown above each plot spectrum. The colour range of each topography is set to vary between plus and minus the maximum absolute value of the data plotted. **A:** The group-level average of the first-level linear trend regressor: the group average GLM t-spectrum (top) and model-predicted magnitude spectra averaged across all sensors for the extrema of the regressors (bottom). **B:** The group-level average of the first-level bad segments regressor, layout is the same as in A. **C:** The group-level average of the first-level V-EOG regressor, layout is the same as in A. **D:** The group-level average of the first-level H-EOG regressor, layout is the same as in A.

Finally, we computed the *Mean* group-level contrast (Contrast 1; Figure 6B) computed on the *V-EOG* and *H-EOG* first-level contrasts separately (Contrast 7 and 8; Figure 1). Three clusters indicated that increased variance in the V-EOG is associated with both increases and decreases in magnitude of the EEG (Figure 8C) One positive effect impacted all sensors and all frequencies (though not always simultaneously) showing the wide-ranging impact of eye movements on the spectrum. Two clusters showed negative effects in the alpha range (around 9Hz) and its harmonic (18Hz) in posterior sensors. This effect indicates a decrease in alpha when eye movements are largest. Increased variance in H-EOG time series was associated with increased magnitude across nearly all sensors and frequencies, as summarised in a single, positive statistical cluster (Figure 8D).

These effects demonstrate that the first-level covariates are associated with consistent group-level effects. However, the results must be interpreted in the context of the variability in first-level effect sizes (Figure 5). Each first-level covariate effect was highly variable in the individual datasets, ranging from no effect in some participants to a covariate effect that exceeded the eyes-open > eyes-closed difference in others (Figure 5B and C). As such, we might not expect to see one or more of these effects in a given single dataset, though the group effects are strong.

### 3.7 Group-level covariate effects

The results so far have combined and contrasted group averages of the first-level results. One possible group-level confound for the age contrast is that the LEMON dataset contains a different number of male and female participants which are not perfectly balanced across age groups. A separate group regressor indicating the reported sex of each participant was included to model between-subject variability relating to this factor, effectively partialling it out from the main age effect of interest in each group contrast. We visualise the overall effect of participant-reported sex on the first-level cope-spectra (averaged across eyes-open and eyes-closed resting-state); this corresponds to the *Sex* group-level contrast (Contrast 5; Figure 6B) computed on the *Overall Mean* first-level contrast (Contrast 1; Figure 1). A single significant cluster identified stronger spectral magnitude in female participants between 1 and 48 Hz, peaking around the alpha and beta frequency ranges (Figure 9A). Increased power in females relative to males has been previously reported in the EEG literature (Aurlien et al., 2004; Zibrandtsen and Kjaer, 2021). Further work is required to distinguish whether this is a true neuronal difference or reflects simpler anatomical differences such as skull thickness.

**Figure 9.**
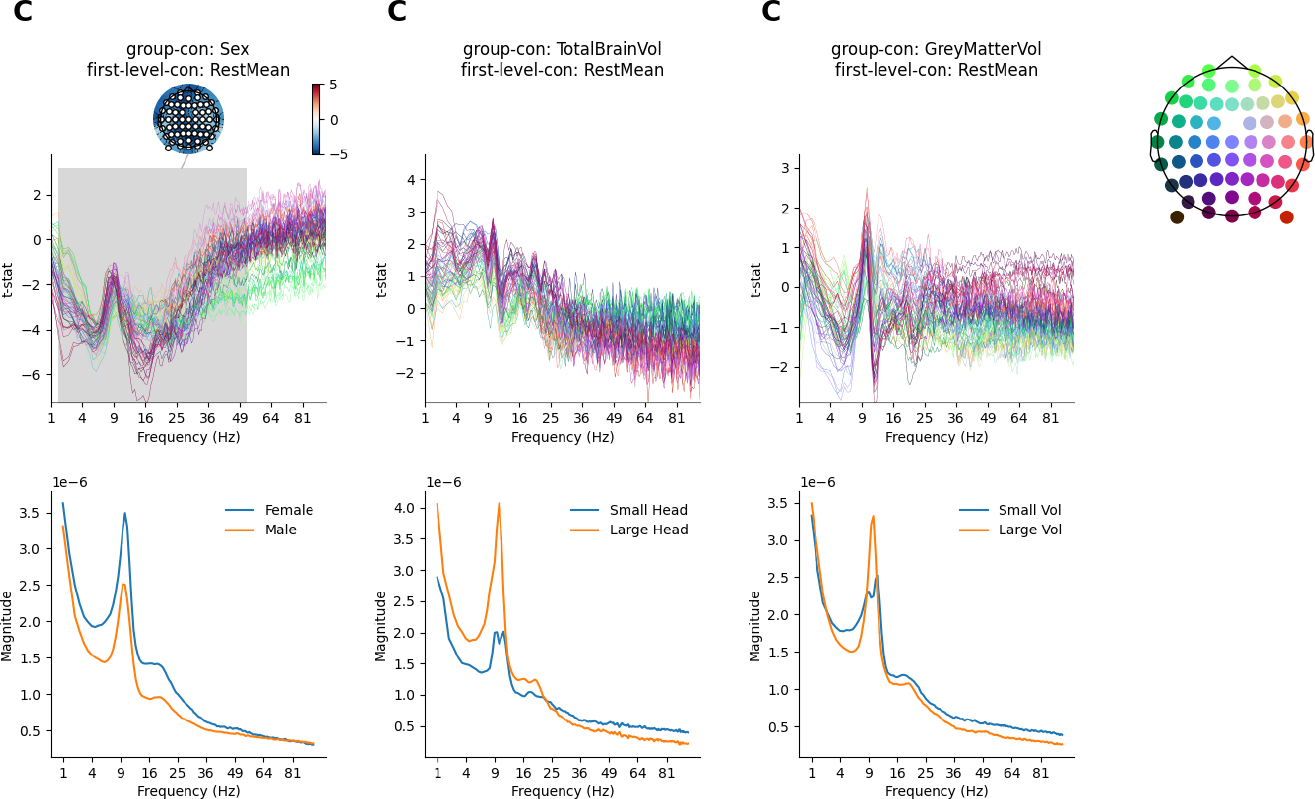
Between-subject group covariate effects on the overall first-level mean. The topography (top right) provides location-colour coding used in the t-spectra throughout the figure. In all cases, statistical significance is assessed by sensors x frequency cluster permutation testing and is indicated in grey. The colour range of each topography is set range between plus and minus the maximum absolute value of the data plotted. **A:** The between-subject difference effect of sex on the average magnitude between females and males. The model projected spectra visualise the group differences (bottom). **B:** Same as A for total brain volume. The model projected spectra visualise the spectrum at the smallest and largest head size. **C:** Same as A for normalised grey matter volume. The model projected spectra visualise the spectrum at the smallest and largest grey matter volume.

Between subject variability associated with two anatomical covariates was modelled at the group-level. The total brain volume of each participant and the proportion of grey matter relative to total brain volume. As before, the inclusion of these regressors ensures that the reported group effects are not biased by these anatomical factors. We separately computed the *TotalBrainVol* and *GrayMatterVol* group-level contrasts (Contrasts 6 and 7; Figure 6B) on the *Overall Mean* first-level contrast (Contrast 1; Figure 1). Non-parametric permutation testing did not identify any significant effects for either overall brain volume or relative grey matter volume (Figure 9B and C). Though these are null effects, the inclusion of these regressors in the group model enables a more refined interpretation of the other results. The inclusion of these confounds means that any variance they might explain can not be attributed to one of the other regressors. Specifically, this increases our confidence that the differences between younger and older adults are not caused by correlated variability in head size or relative grey matter volume.

## 4 Discussion

We have outlined the theory behind the GLM-Spectrum and provided a tutorial overview of its application. We illustrated a practical application for the use of the GLM-Spectrum, using an open EEG dataset to simultaneously quantify and contrast the spectrum of two alternating resting-state conditions whilst regressing out the effect of artefacts including bad segments and eye movements. Both artefact types were associated with a strong group effect but diverse effects at the first-level Of particular interest is the alpha peak in the spectrum of regression coefficients of the V-EOG regressor. This is likely a true neuronal effect linked to blinking that is not removed by ICA but can be explicitly modelled by the GLM-Spectrum. Finally, we extended our analysis to the group-level and explored the spectral differences between older and younger adults, whilst accounting for the effects of sex, brain volume and relative grey matter volume Older adults have lower magnitude in the theta range (3-7 Hz) and higher magnitude in the beta and gamma ranges (>15 Hz). A range of within- and between-subject effects were explored and, crucially, we showed that the reported age effect is robust to differences in participant sex, head size or relative grey matter volume.

### 4.1 A comprehensive framework for spectrum analysis

The GLM-Spectrum is a practical combination of two well-established methodologies that modernises the statistics underlying the time-averaged periodogram, a long-standing and standard spectral estimation method (Bartlett, 1948, 1950; Welch, 1967). Specifically, we utilise multi-level general linear modelling (Friston, 2007; Woolrich et al., 2004), non-parametric permutation testing (Nichols and Holmes, 2001; Winkler et al., 2014), contrast coding and confound regression to extend the scope of classical time-averaged spectrum estimators.

This approach is generalisable to a huge range of analyses. In principle, the GLM-Spectrum could be used in place of Welch’s periodogram or other time-averaged spectrum estimate in any analysis pipeline. A very simple GLM-Spectrum analysis could be configured to be exactly equivalent to these standard approaches. In the simplest case, without first-level covariates the GLM-Spectrum provides a formal framework for multivariate whole head group analysis of power spectra. Moreover, GLM-Spectrum allows for linear denoising of spectrum estimates in datasets where simultaneous recordings of potential artefact sources are available, or artefact time courses can be derived. In addition, covariate effects and contrasts can be readily defined to quickly compute spectra associated with specific external dynamics. For example, an early application of this method has used a GLM-Spectrum to compute power spectra associated with dynamic whole-brain functional networks in MEG (Gohil et al., 2022).

### 4.2 Covariate and confound regression for spectrum analysis

The GLM-Spectrum can characterise spectral changes associated with covariates and potential artefact sources. Standard ICA denoising removes artefacts that share the time-course of the artefact channel. In contrast, confound regression is exploratory across the spectrum. Denoising can be applied to any frequency band with dynamics associated with the segmented artefact time-course irrespective of the original spectrum. For example, the V-EOG blink artefact has a classic low-frequency response that can be attenuated by removing correlated independent components. However, eye blinks are also associated with relatively prolonged changes in alpha and beta power (Liu et al., 2020). In the context of this paper, we consider this to be an ‘indirect’ artefact; it is spatially and spectrally separated from the artefact source and is unlikely to have arisen from volume conduction. The GLM-Spectrum can detect these differences and remove their effect from the overall mean. As such, it could not have been detected or removed by ICA denoising. In another context, this might form the contrast of interest, but in this case, we apply confound regression to minimise the effect of eye movements and blinks on the eyes-open > eyes-closed condition contrast.

### 4.3 Limitations of the GLM-Spectrum model

As outlined in the main text, the parameters of a model are only valid if the underlying assumptions are met. The GLM has several relevant assumptions for the spectrum analysis presented here, particularly at the first-level. In detail, the GLM assumes that the residuals of the model are independently and identically distributed. The presence of any temporal autocorrelation in the residuals indicates that the parameter estimates must be interpreted with caution as this assumption has been violated. Future work can account for this shortcoming by building on similar work in fMRI.

The covariate and confound regressors in a GLM-Spectrum model dynamics over time in a highly simplified sense. This approach is appropriate to quantify relatively slow dynamics, on timescales of seconds, in the context of a spectrum estimator that already utilises sliding time segments for spectrum estimation. The sliding windows are tuned for spectral resolution. They have fixed and arbitrary length and may not accurately reflect the true timescale of dynamics in the covariate variables. As such, limited conclusions about underlying dynamics can be made from a GLM-Spectrum.

We can only say that a dynamic relationship existed at the specific timescale selected for spectrum estimation. If precise temporal dynamics are of interest, a more advanced, window-free method such as the Hidden Markov Model (Quinn et al., 2019, 2018; Vidaurre et al., 2018, 2016) or Empirical Mode Decomposition (Huang et al., 1998) might be more appropriate.

### 4.4 Conclusion

The GLM-Spectrum builds on methodologies that are all well established in the field. The novelty of this work is to bring modern statistics and classical spectrum estimation together into a single framework and to thoroughly explore the theoretical, computational, and practical challenges of its use. The result is an approach for spectrum analysis across the whole head and frequency range with the flexibility to generalise to a huge variety of research and engineering questions.

## 5 Funding & Acknowledgments

This project was supported by the Medical Research Council (RG94383/RG89702) and by the NIHR Oxford Health Biomedical Research Centre. The Wellcome Centre for Integrative Neuroimaging is supported by core funding from the Wellcome Trust (203139/Z/16/Z). A.C.N. is supported by the Wellcome Trust (104571/Z/14/Z) and James S. McDonnell Foundation (220020448). M.W.W. is supported by the Wellcome Trust (106183/Z/14/Z; 215573/Z/19/Z). A.C.N. and M.W.W. are further supported by an EU European Training Network grant (euSSN; 860563). This research was funded in whole, or in part, by the Wellcome Trust . For the purpose of open access, the author has applied a CC BY public copyright licence to any Author Accepted Manuscript version arising from this submission.

The computations described in this paper were performed using the University of Birmingham’s BlueBEAR HPC service, which provides a High Performance Computing service to the University’s research community. See http://www.birmingham.ac.uk/bear for more details.

## 6 Author Contributions

AQ: Conceptualization, Methodology, Software, Formal analysis, Data Curation, Project administration, Writing - Original Draft, Writing - Review & Editing, Visualization. LA: Conceptualization, Methodology, Writing - Review & Editing. CG: Conceptualization, Methodology, Writing - Review & Editing. OK: Methodology, Writing - Review & Editing. JP: Formal analysis, Writing - Review & Editing. CZ: Formal analysis, Writing - Review & Editing. ACN: Writing - Review & Editing, Supervision. MWW: Conceptualization, Methodology, Writing - Original Draft, Writing - Review & Editing, Supervision.

## 7 Supporting information

### 7.1 Time-Averaged Periodogram Estimation

We start by reviewing the definition of the established method for windowed periodogram estimation. The discrete Fourier transform (DFT) can be used to map a series of data points from the time domain into the frequency domain. The frequency domain representation is known as a spectrum and describes how the variance in the data is distributed across frequencies according to a linear basis set. The DFT computes the frequency spectrum *Y* (*f*) from an input time series of real values at discrete time points *y*(*t*).

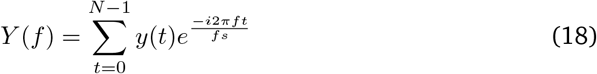

Where *t* is a discrete time point, *f* is a discrete frequency, *fs* is the sampling frequency in Hz and *N* is the number of data points. The output, *Y* (*f*), is a complex-valued array containing the estimate of the spectrum. In practice, the computationally efficient Fast Fourier Transform (FFT; Cooley and Tukey (1965)) implementation of the DFT is applied by most software packages. We will use the FFT for this section, as it refers to the algorithm that is most commonly used in practice.

The mathematics underlying the FFT works with an infinite time series. However, the data from real measurements are finite. Therefore, the FFT must implicitly assume that the time-limited input data, *y*(*t*), repeats infinitely many times. The combination of this repetition and the discrete sampling of the time-series *y*(*t*) leads to the frequency output of the FFT being a linearly spaced axis of N frequency values spanning between 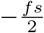 to 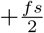. In addition, the discontinuities between repetitions lead to an effect known as spectral leakage, which spreads power contained in one frequency bin to its neighbours. Finally, the computational efficiency of the FFT relies on the input data length *N* being an integer power of 2. If it is not, then the routine will zero-pad the length of the input up to a power of 2. This padding causes further sharp changes and discontinuities that can lead to spectral leakage.

The impact of the spectral leakage is reduced by applying a tapered window function designed to flatten the data at the start and end of each segment to minimise discontinuities between repetitions. Here, we modify equation 18 to multiply a window function *w*(*t*) with the data (point-by-point) during the FFT:

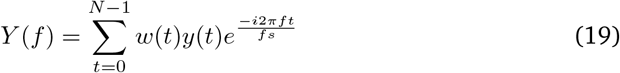

There is a huge range of possible window functions that provide different profiles of spectral leakage and sensitivity. The Hamming and Hann windows (used by MatLab’s ‘pwelch’ function^9^ and SciPy’s ‘scipy.signal.welch’ function ^10^, respectively) are two commonly applied options which offer reasonable narrowband resolution. More advanced tapering can be applied using discrete prolate spheroid sequences (DPSS) to create a Multi-Tapered Spectrum estimate (Prerau et al., 2017; Thomson, 1982).

Y(f) is complex-valued, with its real and imaginary parts reflecting the sine and cosine components of the Fourier transform. The phase and magnitude of each frequency component can be computed from these complex values, though spectrum analyses most commonly use the magnitude. We can take the absolute value of *Y* (*f*) to create a *magnitude spectrum S*_*y*_ (*f*).

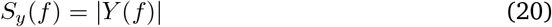

Similarly, a power spectral density (sometimes just called a power spectrum) can be calculated by taking the squared absolute value normalised by the data length:

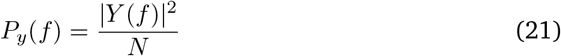

*P*_*y*_ (*f*) is a real-valued estimate of the spectral density of a signal, sometimes known as a periodogram. This periodogram provides a relatively simple spectrum estimation but has a further shortcoming. Only one estimate for the power at each frequency is calculated, irrespective of the length of *y*(*t*). This means that we have no information about the variance around that estimate, and the estimate does not improve if we include more data.

A better estimator is the Time-Averaged Periodogram, introduced by Maurice Bartlett (Bartlett, 1948, 1950) and refined by Peter Welch (Welch, 1967). These methods are commonly used and ensure that the noise level in the periodogram reduces as the length of the input data increases by a method of ensemble averaging (at the expense of information about low frequencies). This splits the input into a set of *k* = 1, 2, … *K* segments (i.e., time windows) each containing *t* = 1, 2, … *T* samples and computes the FFT of each to produce a short time Fourier transform (STFT):

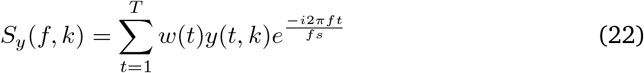

The input for each FFT is now the k-th segment of the continuous input *y*(*t*), which we denote with *y*(*t, k*). The output matrix *Y* (*f, k*) contains the STFT, which describes how the spectrum changes in power across the *K* segments. A time-varying magnitude spectrum can be computed by taking the absolute value of the STFT.

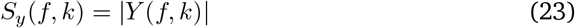

Similarly, a time-varying power spectral density is computed from the squared absolute of the STFT.

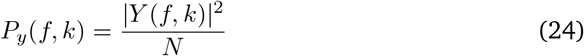

Finally, the time-averaged periodogram is then the average of the time-varying power spectral density across segments. If the previous computations included the windowing function *w*(*t*) and overlapping time segments, then this is Welch’s power spectral density estimate (Welch, 1967).

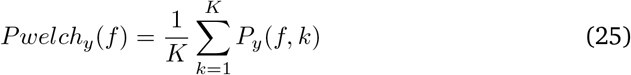

Welch’s time-averaged periodogram now has the property that the noise level of the estimate decreases with increased data length, since more input data provides a larger number of segments for the central averaging step. It is still an imperfect estimator that has been subject to criticism (Prerau et al., 2017; Thomson, 1982) but it is practical, straightforward to compute, and in wide use across science and engineering.

The real-valued power spectrum may additionally be scaled by a log-transform to produce a log-power spectrum *log*(*P*_*y*_ (*f, k*)). This can be desirable as the power spectrum is strictly positive and tends to have a strongly non-Gaussian distribution, whereas the log-power spectrum is not strictly positive and tends to have a more Gaussian distribution (See supplemental section 7.2).

Several key parameters must be set by the user when computing a spectrum in this way. These values affect the range and resolution of the spectrum and must be chosen with care. Briefly, there are three main considerations. Firstly, longer segment lengths (*T*) will increase the frequency resolution (number of bins per Hertz). Secondly, faster data sampling rates allows higher frequencies to be estimated (by increasing the Nyquist range). Finally, increasing the length of the input time-series for a given segment length will increase the number of segments in the average, reducing the impact of noise. These choices are discussed in full in Supplemental section 7.3.

### 7.2 STFT Data Distributions

The general linear model used here expects the dependent variable (the time-varying spectrum) and residuals to follow a Gaussian distribution. Whilst small deviations are permissible, the model fit and any subsequent statistics may be invalid if either is strongly non-Gaussian. This is a concern for the GLM-Spectrum as power values in a standard PSD tend to be non-Gaussian; they are strictly positive and squared (see equation 24). Figure 10A shows an example power spectrum for a single example subject. The second column shows a strongly skewed distribution of power values over time segments for a single channel and frequency. This skew persists into the distribution of residuals as well, indicating that the model assumptions are likely to be violated for when using PSD estimates as the dependent variable in a GLM.

**Figure 10.**
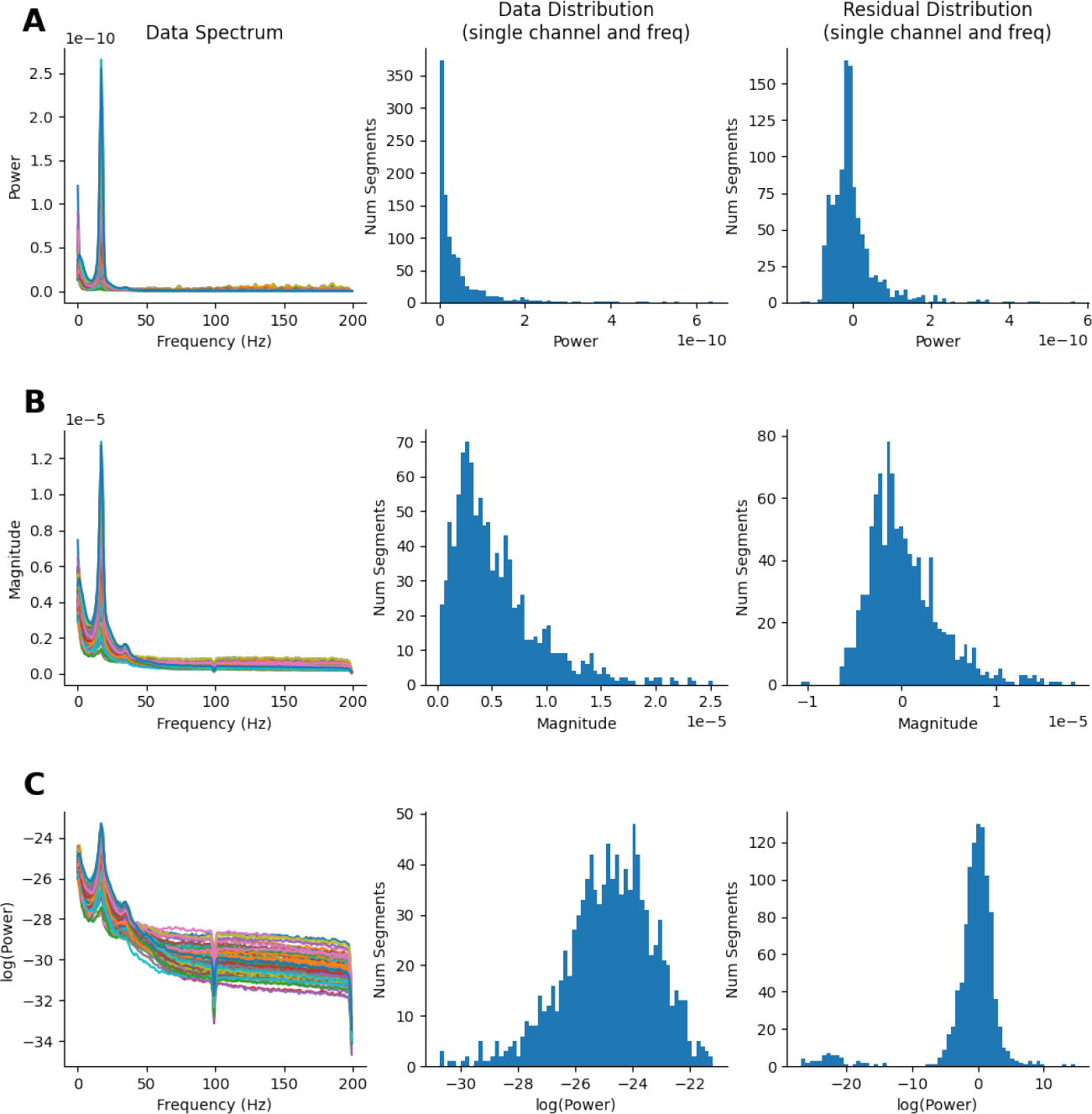
Distribution of the magnitude, power and log-power spectra for an example dataset. **A:** The power spectrum across frequency and channels, the distribution of power values over time segments and the distribution of residuals after fitting a GLM-Spectrum **B:** As A, for the magnitude spectrum. The data and residual distributions are substantially more Gaussian in shape. **C:** As A for the log-power spectrum. The data and residual distributions are more Gaussian than the power and magnitude spectra.

In this paper, we fix this violation by using the magnitude spectrum rather than the power spectrum. Whilst the magnitude spectrum is less commonly used and has less clear mathematical properties ^11^, though the distribution of magnitude estimates is more Gaussian than for power estimates. Figure 10B shows an example magnitude spectrum, its data distribution and its residual distribution. All are better distributed than the PSD example in Figure 10A. Another option would have been to use the log-power spectrum, which also has a well-behaved Gaussian distribution Figure 10C.

Here, we restored the GLM model assumptions through a data-transform however a more general solution could be to use a model that is robust to different data and residual distributions. Future work could explore using a Generalised Linear Model which can describe skewed power distributions with an appropriate link function (Nelder and Wedderburn, 1972).

### 7.3 Parameter settings and resolution in periodograms

When estimating power spectra with a time-averaged approach such as Welch’s Method, three parameters are of particular interest: the sampling rate, the length of the window in which the data will be divided, and the length of the data. Changing these parameters can affect the resolution of the spectrum and how many segments are included in the average.

#### Segment Length

The length of the time segments that the data is divided into influences the resolution of the underlying FFT result. In general, more frequencies are estimated from longer time segments. So, increasing window length whilst holding other parameters constant will result in a larger number of frequency bins in the final spectrum estimate. The resolution in Hz can be computed by dividing the segment length by the sample rate of the data

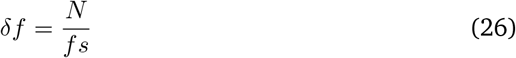

For example, a time series with 512 data points sampled at 128Hz would have a frequency resolution of 4Hz. Similarly, the lowest frequencies that can be reliably extracted from the time-series are also dependent of the window length. Frequencies that are slower than the window period cannot be extracted. In general, it is recommended to estimate power of an oscillatory signal across several cycles to reduce the signal-to-noise ratio of the estimation (Cohen, 2014). Therefore, increasing the window size will assess lower frequencies more reliably (Figure 11A).

**Figure 11.**
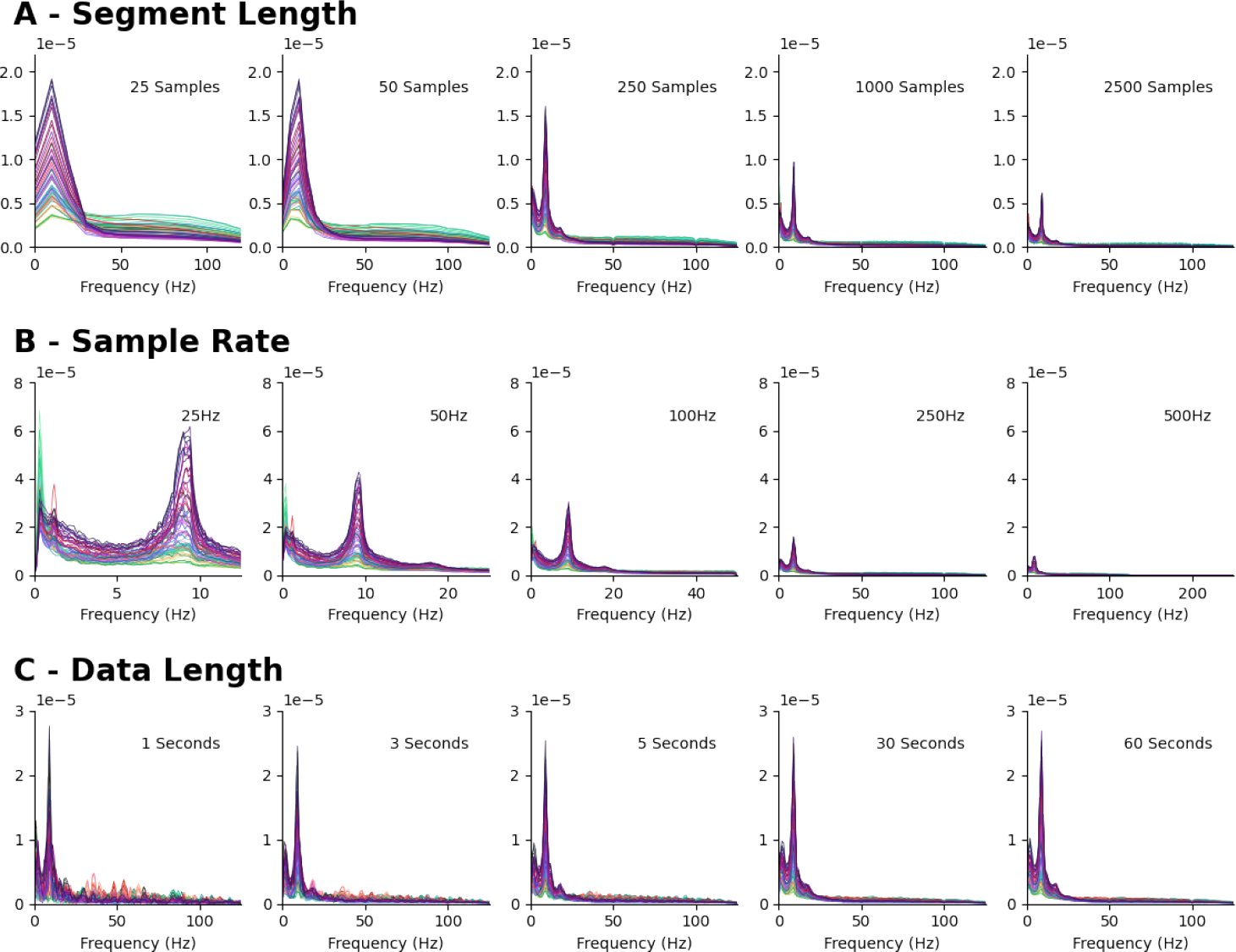
Influence of parameter choice on time-averaged magnitude spectrum estimates. **A:** The effect of increasing segment length on the spectrum estimate. Longer segment lengths have a higher frequency resolution. **B:** The effect of increasing sample rate on the spectrum estimate. Faster sample rates have higher Nyquist frequencies and lower frequency resolution. **C:** The effect on increasing the length of the analysed data set. Longer datasets have more segments contributing to the central average and are less affected by noise.

There is an additional subtlety in some implementations of the FFT that means that the data length does not necessarily equal the length of the computed FFT. For example, the scipy implementation of Welch’s method contains parameters for segment length (nperseg) and FFT length (nfft). These are assigned equivalent values by default, but if they are different to the nfft parameter should be used instead of N to compute the frequency resolution.

#### Sampling Rate

The FFT returns estimates at equally spaced frequencies from 0Hz to one half of the sampling rate of the data Nyquist Frequency. The 0Hz component is known as a ‘direct current’ or DC offset and contains the average of all samples in the segment being analysed. The Nyquist frequency is the fastest observable frequency in the dataset and is one half of the sample rate. As such, increases in the sampling rate, whilst holding other parameters constant, will in result in larger frequency spacing and a lower resolution (Figure 11A). The reason for this is that the same number of frequency bins must cover a larger frequency range. In reverse, this means that decreasing the sampling rate while holding the window length constant results in a smaller frequency spacing and higher spectral resolution.

Importantly, increasing the frequency spacing is not always better because smaller bins might result in a noisier estimation of the power spectrum (Figure 11B). Small variations in the (intrinsic) frequency of the oscillatory signals - that are in most cases not of particular interest - might be represented by power in various of these small frequency bins whereas larger frequency bins smooth across these smaller, less relevant variations. This results in a smoother power spectrum.

#### Length of the overall time-series

The length of the data determines the number of windows in which the time course is divided by Welch’s Method before averaging across power spectra. A higher number of windows results in averaging across a larger number of power spectra which in turn results in a better signal-to-noise ratio of the overall power estimation (Figure 11C). In general, considering more data is preferable because the average of a larger number of window-power spectra should result in an estimation closer to the true average power spectrum according to the law of large numbers. Importantly, dynamic fluctuations in oscillatory power over the time course might introduce meaningful variations in the mean power spectrum that cannot be accounted for by Welch’s Method and result in more complex, and less peaky power spectra.

### 7.4 Fast standard-error estimation

The slowest computation in the GLM-Spectrum is obtaining the standard error of the parameter estimates, known as varcopes. This involves large matrix multiplications that are typically repeated across many tests. However, only the diagonal output is used in the eventual varcope estimate. Standard computing approaches will evaluate every single cell within these matrices even though majority contribute to the off-diagonals and are eventually discarded. This is not a large computational expense for single tests but quickly becomes prohibitive when computing large numbers of test together in matrix form. We use the numpy implementation of Einstein summation conventions (Numpy.Einsum — NumPy v1.23 Manual, n.d.)(https://numpy.org/doc/stable/reference/generated/numpy.einsum.html) to compute the most efficient computational path for the varcope evaluation. In practice, this avoids carrying out intermediate multiplications which would eventually be dropped when extracting the diagonal at the final step. The code for the standard ‘DotDiag’ approach and the new Einsum approache is as follows:

We explore the difference this makes to computation time by running 100 simulated GLMs with each method for each of four datasets. The first dataset was a single GLM that might represent a single channel and frequency bin. The second is a full GLM-Spectrum across 100 frequency bins whilst the third and fourth were GLM-Spectra across 60 or 204 channels, representing common EEG and MEG data sizes. The GLM was computed 100 times for each case and the results are summarised in Figure 12. The computation was equally fast for both methods in the single-test case but the einsum approach can lead to a 1000x speed up in computation time in the larger datasets. This is convenient for in any case but is essential for making non-parametric permutation statistics of GLM-spectra practically feasible.

**Figure 12.**
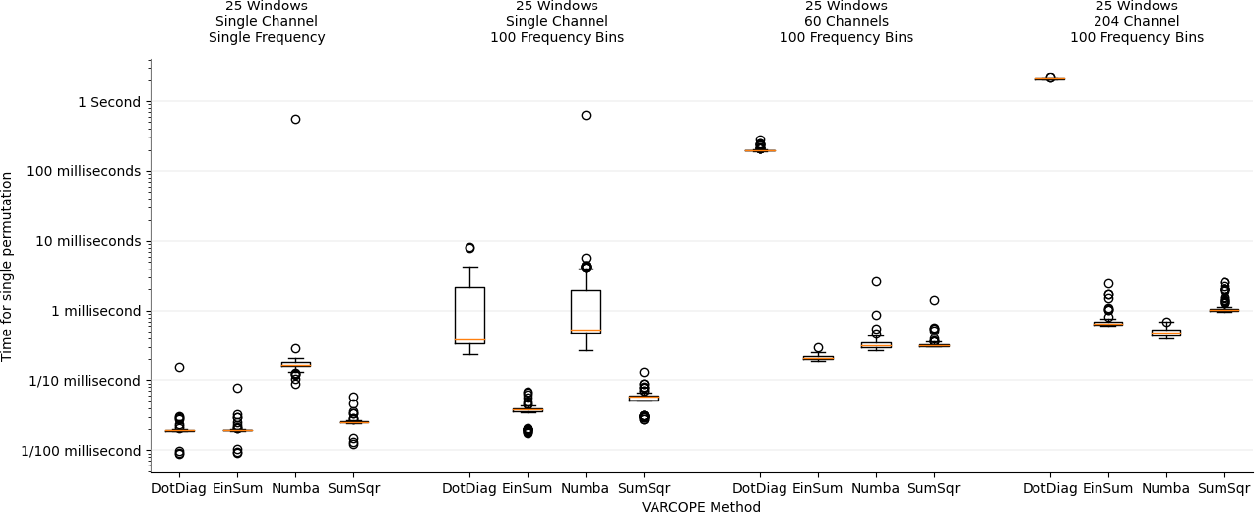
Computation times for varcope estimation across different data sizes with a standard or einsum based method. Each data size and method was computed for random data 100 times and the distribution of its timing shows as a boxplot.

**Figure 13.**
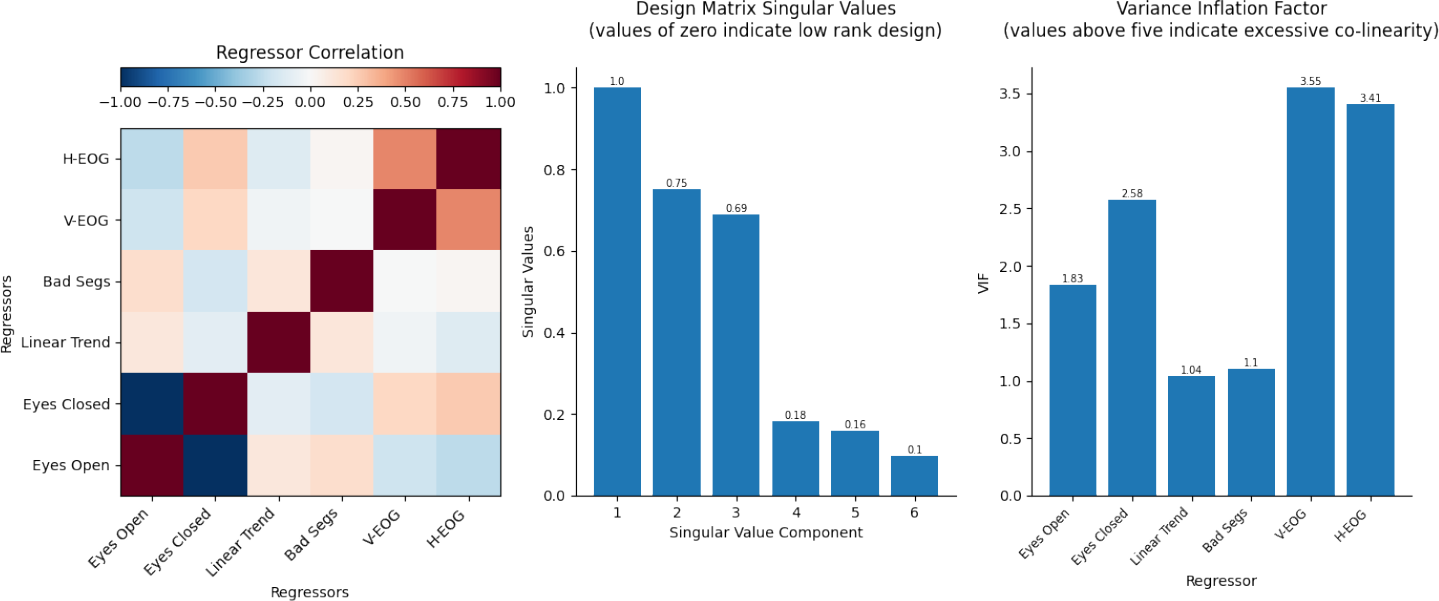
The regressor correlations, singular-value spectrum and variance inflation factors for the first-level GLM-Spectrum design matrix for a single dataset. This analysis indicates that there is some indication of co-linearity but not at a problematic level as the smallest singular-values do not reach zero, and the variance inflation factors do not exceed 5.

**Figure 14.**
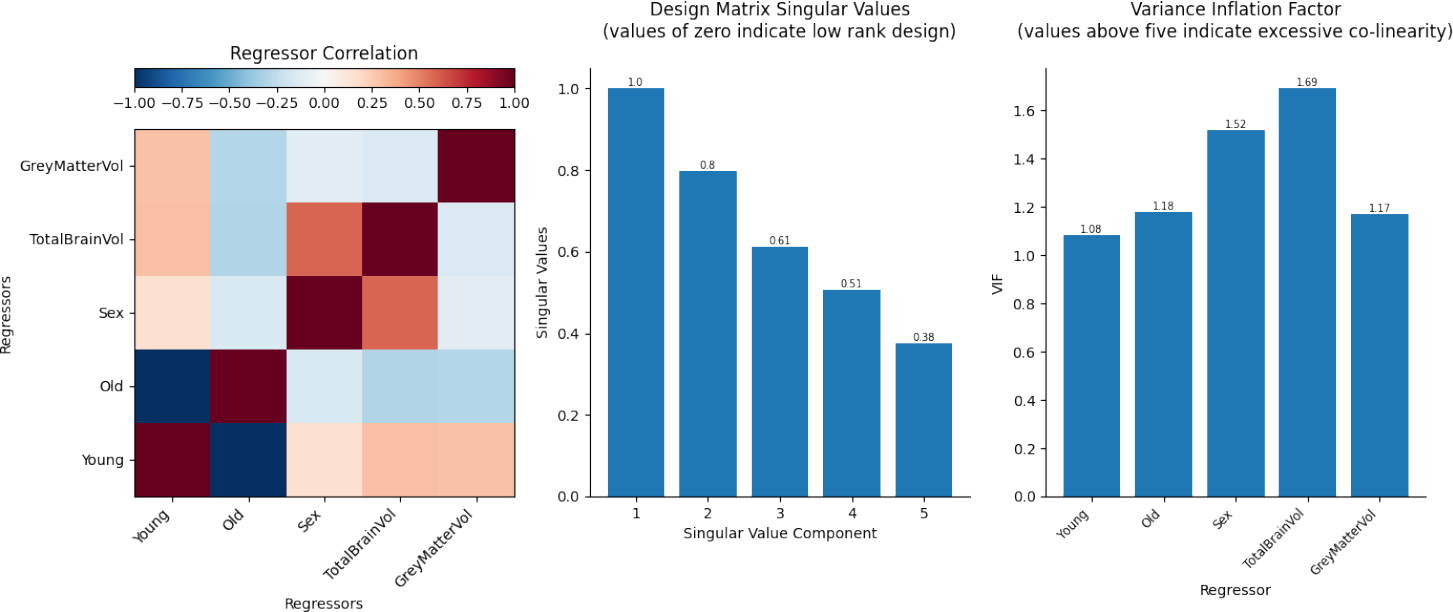
The regressor correlations, singular-value spectrum and variance inflation factors for the group-level GLM-Spectrum design matrix. This analysis indicates that there is some indication of co-linearity but not at a problematic level as the smallest singular-values do not reach zero, and the variance inflation factors do not exceed 5.

### 7.5 GLM Design Efficiency

### 7.6 Confound Regression

Figure 15A shows an example design matrix for this case. The first regressor is a constant vector of ones and a second regressor tracks the time segments in which the artefact occurs. In isolation, the constant regressor models the data mean (including the time periods that the artefact occurs), but this changes when it is alongside a regressor with non-zero mean in the same model. The constant regressor now models the intercept; this is the expected value of the data when the artefact regressor is zero (Figure 15B – contrast 1 ‘Intercept’). In turn, the artefact regressor can be interpreted as the difference between the value of the intercept and the mean of the segments indicated in the artefact regressor (Figure 15B – contrast 2 ‘Artefact Effect’). We can recover the mean of the artefact segments by summing the parameter estimates for both regressors (Figure 15B – contrast 3 ‘Artefact Mean’).

**Figure 15.**
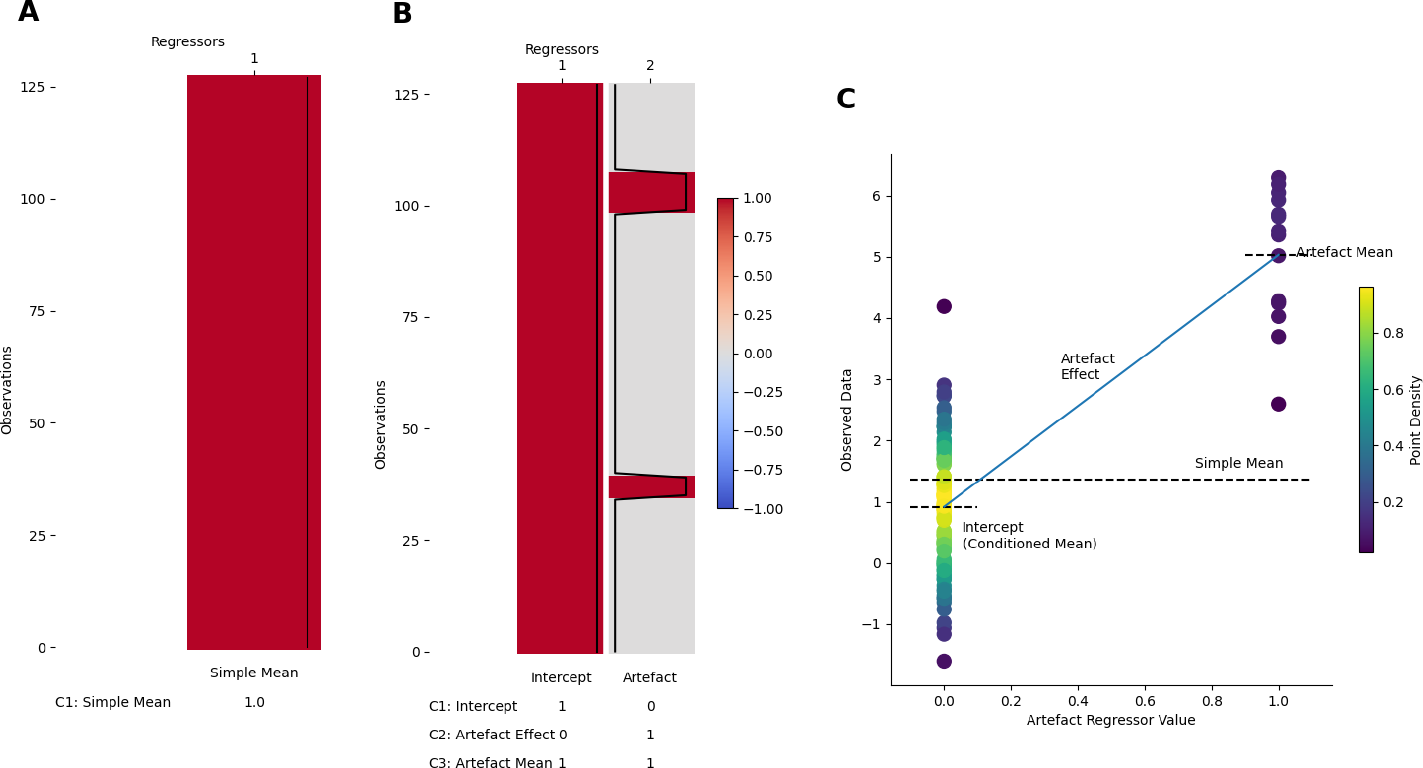
GLM-Spectrum design matrix for confound regression. **A:** A GLM design with a single regressor and contrast modelling a simple mean of the observed data **B:** A GLM design with two regressors and three contrasts illustrating confound regression. The first regressor is a constant term and the second is a sparse, non-zero mean artefact regressor indicating data observations that are possible artefacts. Three contrasts isolate each regressor individually (intercept and artefact effect) and their sum (artefact mean). **C:** Simulated data observations and the quantities estimated by the GLM designs in A and B. The first design estimates the simple mean overall data, though this is heavily influenced by the possible outlier points. The second design models an intercept and an artefact effect. The intercept can be thought of as the mean where the artefact regressor equals zero, and the artefact effect is the distance between the intercept and the artefact mean. Finally, the absolute artefact mean can be reconstructed by the sum of the two parameter estimates as shown in B contrast 3.

To illustrate the quantities estimated by the simple model and the confound regression, we generate a simulation of 128 data points centred around a ‘true’ mean of 1 with a small number of outliers centred around 4 (Figure 15C). The simple mean estimated by the ‘mean-only’ design is biased towards the outlier observations. In contrast, the confound regression quantifies the ‘true mean’ of 1 in the intercept term as the artefact regressor describes the effect of the artefact. In a real data example, this design could describe the mean spectrum of a resting-state EEG recording whilst accounting for a set of ‘bad segments’ annotations identified during pre-processing. This both provides an estimate of any ‘artefact effect’ and linearly removes its influence from the estimate of the mean term.

We can also specify artefact regressor with the mean removed, i.e. with zero means. Counter-intuitively, the interpretation of the regression parameter estimate is unchanged; whereas the interpretation of the constant regressor changes from modelling the intercept (as in Figure 15B) to modelling the mean over all time points.

### 7.7 Effect of Confound Regression on Eyes-Open < Eyes-Closed Contrast

Adding covariate and confound regressors to a model can affect the other parameter estimates and contrasts. This effect is particularly strong when there are correlations between any of the new and existing regressors. For example, the current resting-state recordings interleaves periods of eyes-open and eyes close rest so it is likely that eye movements will increase in the eyes-open periods. This correlation means that a some eye movement related effects might leak into contrast between the eyes-open and eyes-closed conditions unless they are explicitly modelled.

Here, we illustrate this with the group-level average t-spectra for the first-level eyes-open> eyes-closed contrast. This is shown for two different first-level models; a reduced model with no eye movement covariates and a full model with first-level confound modelling. There is a large difference in low frequencies in frontal sensors though the t-spectra are similar in posterior and occipital sensors. Without first-level confound modelling, a large frontal effect can be seen between 1Hz and 4Hz with t-values peaking around 5 (Figure 16A). This is strongly attenuated with the additional confound modelling at the first-level (Figure 16B) which reduces the open > closed contrast t-values in frontal, low frequency regions by around 4.

**Figure 16.**
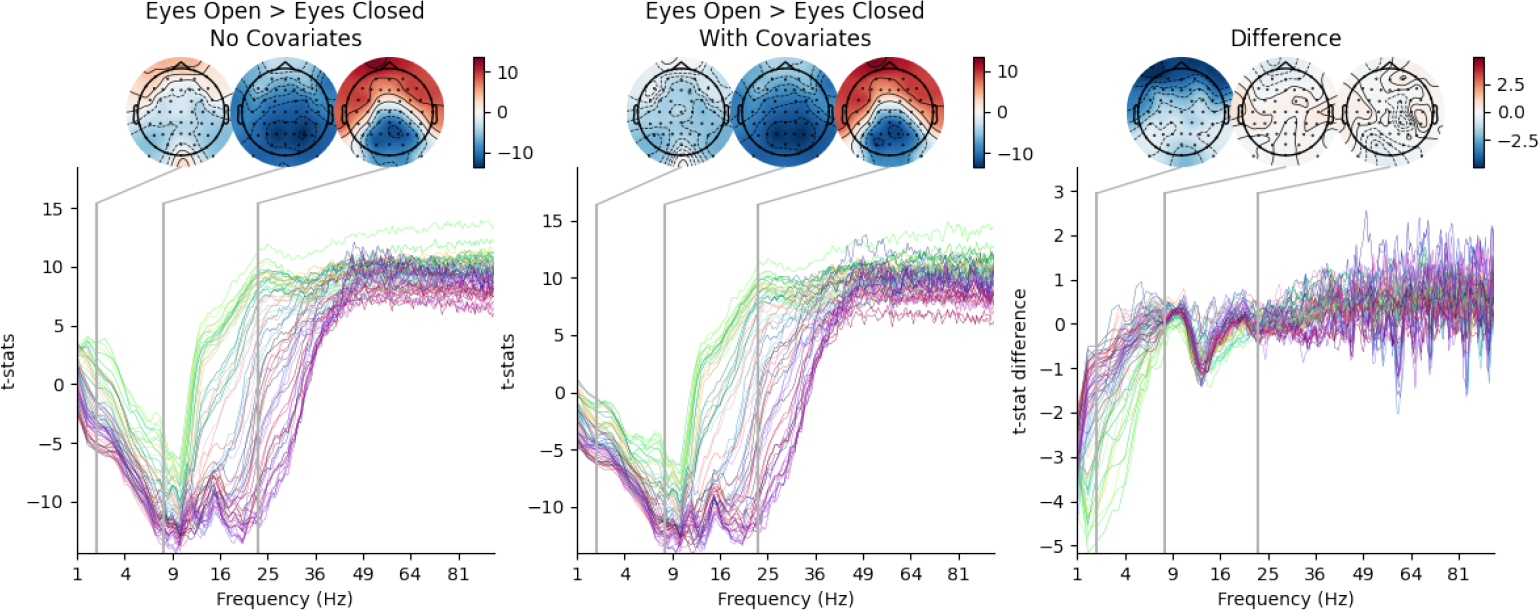
Effect of first-level confound regressors on group-level eyes-open > eyes-closed contrast. **A:** A t-spectrum of the contrast between eyes-open and eyes-closed resting-states with no covariate or confound regressors. **B:** A t-spectrum of the same contrast as A from a GLM with an additional confound regressor modelling V-EOG activity. **C:** The difference between A and B.

## Footnotes

1 The term ‘beta’ has different uses in the fields of statistics/linear-modelling and neuronal oscillations. In this work, we refer to a ‘beta-spectrum’ in the statistical sense (a spectrum of linear model parameter estimates) rather than the neuronal sense (a spectrum of beta-oscillation activity).

2 https://numpy.org/doc/stable/reference/generated/numpy.einsum.html

3 Cohen’s original work suggest that values of *F*^2^ ⩾ 0.02, *F*^2^ ⩾ 0.15 and *F*^2^ ⩾ 0.35 represent small, medium and large effects respectively (Cohen, 1988; Selya et al., 2012). These labels should be interpreted with caution and only used as informal guidelines.

4 https://fsl.fmrib.ox.ac.uk/fsl/fslwiki/GLM

5 https://github.com/OHBA-analysis/osl

6 https://github.com/nbara/python-meegkit

7 https://pypi.org/project/glmtools/

8 https://github.com/OHBA-analysis/Quinn2022_GLMSpectrum

9 https://www.mathworks.com/help/signal/ref/pwelch.html

10 ^10^https://docs.scipy.org/doc/scipy/reference/generated/scipy.signal.welch.html

11 The relation between the sum-square of the time-domain data and the integral of the spectrum does not hold for a magnitude spectrum.

## References

Adrian, E. D. and Matthews, B. H. C. (1934). The berger rhythm: Potential change from the occpital lobes in man. Brain, 57(4):355–385.

Aurlien, H., Gjerde, I. O., Aarseth, J. H., Eldøen, G., Karlsen, B., Skeidsvoll, H., and Gilhus, N. E. (2004). EEG background activity described by a large computerized database. Clinical Neurophysiology, 115(3):665–673.

Babayan, A., Erbey, M., Kumral, D., Reinelt, J. D., Reiter, A. M. F., Röbbig, J., Schaare, H. L., Uhlig, M., Anwander, A., Bazin, P.-L., Horstmann, A., Lampe, L., Nikulin, V. V., Okon-Singer, H., Preusser, S., Pampel, A., Rohr, C. S., Sacher, J., Thöne-Otto, A., Trapp, S., Nierhaus, T., Altmann, D., Arelin, K., Blöchl, M., Bongartz, E., Breig, P., Cesnaite, E., Chen, S., Cozatl, R., Czerwonatis, S., Dambrauskaite, G., Dreyer, M., Enders, J., Engelhardt, M., Fischer, M. M., Forschack, N., Golchert, J., Golz, L., Guran, C. A., Hedrich, S., Hentschel, N., Hoffmann, D. I., Huntenburg, J. M., Jost, R., Kosatschek, A., Kunzendorf, S., Lammers, H., Lauckner, M. E., Mahjoory, K., Kanaan, A. S., Mendes, N., Menger, R., Morino, E., Näthe, K., Neubauer, J., Noyan, H., Oligschläger, S., Panczyszyn-Trzewik, P., Poehlchen, D., Putzke, N., Roski, S., Schaller, M.-C., Schieferbein, A., Schlaak, B., Schmidt, R., Gorgolewski, K. J., Schmidt, H. M., Schrimpf, A., Stasch, S., Voss, M., Wiedemann, A., Margulies, D. S., Gaebler, M., and Villringer, A. (2019). A mind-brain-body dataset of MRI, EEG, cognition, emotion, and peripheral physiology in young and old adults. Scientific Data, 6(1).

Babiloni, C., Lizio, R., Vecchio, F., Frisoni, G. B., Pievani, M., Geroldi, C., Claudia, F., Ferri, R., Lanuzza, B., and Rossini, P. M. (2011). Reactivity of cortical alpha rhythms to eye opening in mild cognitive impairment and alzheimer’s disease: an EEG study. Journal of Alzheimer’s Disease, 22(4):1047–1064.

Baker, D. H. (2021). Statistical analysis of periodic data in neuroscience. Neurons, Behavior, Data analysis, and Theory, 5(3).

Bartlett, M. S. (1948). Smoothing periodograms from time-series with continuous spectra. Nature, 161(4096):686–687.

Bartlett, M. S. (1950). Perioidogram analysis and continuous spectra. Biometrika, 37(1-2):1–16.

Beckmann, C. F., Jenkinson, M., and Smith, S. M. (2003). General multilevel linear modeling for group analysis in FMRI. NeuroImage, 20(2):1052–1063.

Benwell, C. S., London, R. E., Tagliabue, C. F., Veniero, D., Gross, J., Keitel, C., and Thut, G. (2019). Frequency and power of human alpha oscillations drift systematically with time-on-task. NeuroImage, 192:101–114.

Buzsakḱi, G. and Draguhn, A. (2004). Neuronal oscillations in cortical networks. Science, 304(5679):1926–1929.

Cohen, J. (1988). Statistical power analysis for the behavioral sciences. L. Erlbaum Associates, Hillsdale, N.J, 2nd ed edition.

Cohen, M. X. (2014). Analyzing Neural Time Series Data. The MIT Press.

Cooley, J. W. and Tukey, J. W. (1965). An algorithm for the machine calculation of complex fourier series. Mathematics of Computation, 19(90):297–301.

de Cheveigné, A. (2020). ZapLine: A simple and effective method to remove power line artifacts. NeuroImage, 207:116356.

Draper, N. R. and Stoneman, D. M. (1966). Testing for the inclusion of variables in linear regression by a randomisation technique. Technometrics, 8(4):695.

Freedman, D. and Lane, D. (1983). A nonstochastic interpretation of reported significance levels. Journal of Business and Economic Statistics, 1(4):292–298.

Friston, K., Josephs, O., Zarahn, E., Holmes, A., Rouquette, S., and Poline, J.-B. (2000). To smooth or not to smooth? NeuroImage, 12(2):196–208.

Friston, K., Penny, W., Phillips, C., Kiebel, S., Hinton, G., and Ashburner, J. (2002). Classical and bayesian inference in neuroimaging: Theory. NeuroImage, 16(2):465–483.

Friston, K. J., editor (2007). Statistical parametric mapping: the analysis of funtional brain images. Elsevier/Academic Press, Amsterdam ; Boston, 1st ed edition.

Friston, K. J., Holmes, A. P., Worsley, K. J., Poline, J.-P., Frith, C. D., and Frackowiak, R. S. J. (1994). Statistical parametric maps in functional imaging: A general linear approach. Human Brain Mapping, 2(4):189–210.

Gelman, A. and Hill, J. (2007). Data analysis using regression and multilevel/hierarchical models. Analytical methods for social research. Cambridge University Press, Cambridge ; New York. OCLC: ocm67375137.

Gohil, C., Roberts, E., Timms, R., Skates, A., Higgins, C., Quinn, A., Pervaiz, U., van Amersfoort, J., Notin, P., Gal, Y., Adaszewski, S., and Woolrich, M. (2022). Mixtures of large-scale dynamic functional brain network modes. NeuroImage, 263:119595.

Gramfort, A. (2013). MEG and EEG data analysis with MNE-python. Frontiers in Neuroscience, 7.

Greve, D. N. and Fischl, B. (2009). Accurate and robust brain image alignment using boundary-based registration. NeuroImage, 48(1):63–72.

Harris, C. R., Millman, K. J., van der Walt, S. J., Gommers, R., Virtanen, P., Cournapeau, D., Wieser, E., Taylor, J., Berg, S., Smith, N. J., Kern, R., Picus, M., Hoyer, S., van Kerkwijk, M. H., Brett, M., Haldane, A., del Río, J. F., Wiebe, M., Peterson, P., Gérard-Marchant, P., Sheppard, K., Reddy, T., Weckesser, W., Abbasi, H., Gohlke, C., and Oliphant, T. E. (2020). Array programming with NumPy. Nature, 585(7825):357–362.

Huang, N. E., Shen, Z., Long, S. R., Wu, M. C., Shih, H. H., Zheng, Q., Yen, N.-C., Tung, C. C., and Liu, H. H. (1998). The empirical mode decomposition and the hilbert spectrum for nonlinear and non-stationary time series analysis. Proceedings of the Royal Society of London. Series A: Mathematical, Physical and Engineering Sciences, 454(1971):903–995.

Hunter, J. D. (2007). Matplotlib: A 2d graphics environment. Computing in Science and Engineering, 9(3):90–95.

Hyvarinen, A. (1999). Fast and robust fixed-point algorithms for independent component analysis. IEEE Transactions on Neural Networks, 10(3):626–634.

Jenkinson, M., Bannister, P., Brady, M., and Smith, S. (2002). Improved optimization for the robust and accurate linear registration and motion correction of brain images. NeuroImage, 17(2):825–841.

Jenkinson, M. and Smith, S. (2001). A global optimisation method for robust affine registration of brain images. Medical Image Analysis, 5(2):143–156.

Kayser, J. and Tenke, C. E. (2015). On the benefits of using surface laplacian (current source density) methodology in electrophysiology. International Journal of Psychophysiology, 97(3):171–173.

Klimesch, W. (1999). EEG alpha and theta oscillations reflect cognitive and memory performance: a review and analysis. Brain Research Reviews, 29(2-3):169–195.

Knief, U. and Forstmeier, W. (2021). Violating the normality assumption may be the lesser of two evils. Behavior Research Methods, 53(6):2576–2590.

Kopell, N. J., Gritton, H. J., Whittington, M. A., and Kramer, M. A. (2014). Beyond the connectome: The dynome. Neuron, 83(6):1319–1328.

Litvak, V., Jha, A., Flandin, G., and Friston, K. (2013). Convolution models for induced electromagnetic responses. NeuroImage, 64:388–398.

Liu, C. C., Hajra, S. G., Pawlowski, G., Fickling, S. D., Song, X., and D’Arcy, R. C. (2020). Differential neural processing of spontaneous blinking under visual and auditory sensory environments: An EEG investigation of blink-related oscillations. NeuroImage, 218:116879.

Merkin, A., Sghirripa, S., Graetz, L., Smith, A. E., Hordacre, B., Harris, R., Pitcher, J., Semmler, J., Rogasch, N. C., and Goldsworthy, M. (2022). Do age-related differences in aperiodic neural activity explain differences in resting EEG alpha? Neurobiology of Aging.

Nelder, J. A. and Wedderburn, R. W. M. (1972). Generalized linear models. Journal of the Royal Statistical Society. Series A (General), 135(3):370.

Nichols, T. E. and Holmes, A. P. (2001). Nonparametric permutation tests for functional neuroimaging: A primer with examples. Human Brain Mapping, 15(1):1–25.

O’Gorman, T. W. (2005). The performance of randomization tests that use permutations of independent variables. Communications in Statistics - Simulation and Computation, 34(4):895–908.

Penrose, R. (1956). On best approximate solutions of linear matrix equations. Mathematical Proceedings of the Cambridge Philosophical Society, 52(1):17–19.

Perrin, F., Pernier, J., Bertrand, O., and Echallier, J. (1989). Spherical splines for scalp potential and current density mapping. Electroencephalography and Clinical Neurophysiology, 72(2):184–187.

Prerau, M. J., Brown, R. E., Bianchi, M. T., Ellenbogen, J. M., and Purdon, P. L. (2017). Sleep neurophysiological dynamics through the lens of multitaper spectral analysis. Physiology, 32(1):60–92.

Quinn, A. and Hymers, M. (2020). SAILS: Spectral analysis in linear systems. Journal of Open Source Software, 5(47):1982.

Quinn, A. J., van Ede, F., Brookes, M. J., Heideman, S. G., Nowak, M., Seedat, Z. A., Vidaurre, D., Zich, C., Nobre, A. C., and Woolrich, M. W. (2019). Unpacking transient event dynamics in electrophysiological power spectra. Brain Topography, 32(6):1020–1034.

Quinn, A. J., van Es, M., Gohil, C., and Woolrich, M. W. (2022). Ohba software library in python (osl). https://zenodo.org/record/6875060.

Quinn, A. J., Vidaurre, D., Abeysuriya, R., Becker, R., Nobre, A. C., and Woolrich, M. W. (2018). Task-evoked dynamic network analysis through hidden markov modeling. Frontiers in Neuroscience, 12.

Rosner, B. (1983). Percentage points for a generalized ESD many-outlier procedure. Technometrics, 25(2):165–172.

Selya, A. S., Rose, J. S., Dierker, L. C., Hedeker, D., and Mermelstein, R. J. (2012). A practical guide to calculating cohen’s f2, a measure of local effect size, from PROC MIXED. Frontiers in Psychology, 3.

Smith, N. J. and Kutas, M. (2014). Regression-based estimation of ERP waveforms: I. the rERP framework. Psychophysiology, 52(2):157–168.

Smith, S., Jenkinson, M., Beckmann, C., Miller, K., and Woolrich, M. (2007). Meaningful design and contrast estimability in FMRI. NeuroImage, 34(1):127–136.

Smith, S. M. (2002). Fast robust automated brain extraction. Human Brain Mapping, 17(3):143–155.

Thomson, D. (1982). Spectrum estimation and harmonic analysis. Proceedings of the IEEE, 70(9):1055–1096.

Vidaurre, D., Hunt, L. T., Quinn, A. J., Hunt, B. A. E., Brookes, M. J., Nobre, A. C., and Woolrich, M. W. (2018). Spontaneous cortical activity transiently organises into frequency specific phase-coupling networks. Nature Communications, 9(1).

Vidaurre, D., Quinn, A. J., Baker, A. P., Dupret, D., Tejero-Cantero, A., and Woolrich, M. W. (2016). Spectrally resolved fast transient brain states in electrophysiological data. NeuroImage, 126:81–95.

Virtanen, P., Gommers, R., Oliphant, T. E., Haberland, M., Reddy, T., Cournapeau, D., Burovski, E., Peterson, P., Weckesser, W., Bright, J., van der Walt, S. J., Brett, M., Wilson, J., Millman, K. J., Mayorov, N., Nelson, A. R. J., Jones, E., Kern, R., Larson, E., Carey, C. J., Polat, i., Feng, Y., Moore, E. W., VanderPlas, J., Laxalde, D., Perktold, J., Cimrman, R., Henriksen, I., Quintero, E. A., Harris, C. R., Archibald, A. M., Ribeiro, A. H., Pedregosa, F., van Mulbregt, P., Vijaykumar, A., Bardelli, A. P., Rothberg, A., Hilboll, A., Kloeckner, A., Scopatz, A., Lee, A., Rokem, A., Woods, C. N., Fulton, C., Masson, C., Häggström, C., Fitzgerald, C., Nicholson, D. A., Hagen, D. R., Pasechnik, D. V., Olivetti, E., Martin, E., Wieser, E., Silva, F., Lenders, F., Wilhelm, F., Young, G., Price, G. A., Ingold, G.-L., Allen, G. E., Lee, G. R., Audren, H., Probst, I., Dietrich, J. P., Silterra, J., Webber, J. T., Slavič, J., Nothman, J., Buchner, J., Kulick, J., Schönberger, J. L., de Miranda Cardoso, J. V., Reimer, J., Harrington, J., Rodríguez, J. L. C., Nunez-Iglesias, J., Kuczynski, J., Tritz, K., Thoma, M., Newville, M., Kümmerer, M., Bolingbroke, M., Tartre, M., Pak, M., Smith, N. J., Nowaczyk, N., Shebanov, N., Pavlyk, O., Brodtkorb, P. A., Lee, P., McGibbon, R. T., Feldbauer, R., Lewis, S., Tygier, S., Sievert, S., Vigna, S., Peterson, S., More, S., Pudlik, T., Oshima, T., Pingel, T. J., Robitaille, T. P., Spura, T., Jones, T. R., Cera, T., Leslie, T., Zito, T., Krauss, T., Upadhyay, U., Halchenko, Y. O., and and, Y. V.-B. (2020). SciPy 1.0: fundamental algorithms for scientific computing in python. Nature Methods, 17(3):261–272.

Voytek, B., Kramer, M. A., Case, J., Lepage, K. Q., Tempesta, Z. R., Knight, R. T., and Gazzaley, A. (2015). Age-Related Changes in 1/f Neural Electrophysiological Noise. Journal of Neuroscience, 35(38):13257–13265.

Wan, L., Huang, H., Schwab, N., Tanner, J., Rajan, A., Lam, N. B., Zaborszky, L., shan R. Li, C., Price, C. C., and Ding, M. (2018). From eyes-closed to eyes-open: Role of cholinergic projections in EC-to-EO alpha reactivity revealed by combining EEG and MRI. Human Brain Mapping, 40(2):566–577.

Welch, P. (1967). The use of fast fourier transform for the estimation of power spectra: A method based on time averaging over short, modified periodograms. IEEE Transactions on Audio and Electroacoustics, 15(2):70–73.

Williams, M. N., Grajales, C. A. G., and Kurkiewicz, D. (2013). Assumptions of multiple regression: Correcting two misconceptions.

Winkler, A. M., Ridgway, G. R., Webster, M. A., Smith, S. M., and Nichols, T. E. (2014). Permutation inference for the general linear model. NeuroImage, 92:381–397.

Woolrich, M. W., Behrens, T. E., Beckmann, C. F., Jenkinson, M., and Smith, S. M. (2004). Multilevel linear modelling for FMRI group analysis using bayesian inference. NeuroImage, 21(4):1732–1747.

Woolrich, M. W., Jbabdi, S., Patenaude, B., Chappell, M., Makni, S., Behrens, T., Beckmann, C., Jenkinson, M., and Smith, S. M. (2009). Bayesian analysis of neuroimaging data in FSL. NeuroImage, 45(1):S173–S186.

Woolrich, M. W., Ripley, B. D., Brady, M., and Smith, S. M. (2001). Temporal autocorrelation in univariate linear modeling of FMRI data. NeuroImage, 14(6):1370–1386.

Worsley, K. and Friston, K. (1995). Analysis of fMRI time-series revisited—again. NeuroImage, 2(3):173–181.

Xifra-Porxas, A., Niso, G., Larivière, S., Kassinopoulos, M., Baillet, S., Mitsis, G. D., and Boudrias, M.-H. (2019). Older adults exhibit a more pronounced modulation of beta oscillations when performing sustained and dynamic handgrips. NeuroImage, 201:116037.

Zhang, Y., Brady, M., and Smith, S. (2001). Segmentation of brain MR images through a hidden markov random field model and the expectation-maximization algorithm. IEEE Transactions on Medical Imaging, 20(1):45–57.

Zibrandtsen, I. C. and Kjaer, T. W. (2021). Fully automatic peak frequency estimation of the posterior dominant rhythm in a large retrospective hospital EEG cohort. Clinical Neurophysiology Practice, 6:1–9.

